# Endosomes deliver ceramide phosphoethanolamine with unique acyl chain anchors to the cleavage furrow during male meiotic cytokinesis

**DOI:** 10.1101/2021.04.27.441622

**Authors:** Govind Kunduri, Si-Hung Le, Valentina Baena, Nagampalli Vijaykrishna, Adam Harned, Kunio Nagashima, Daniel Blankenberg, Yoshihiro Izumi, Kedar Narayan, Takeshi Bamba, Usha Acharya, Jairaj K. Acharya

## Abstract

Cell division, wherein one cell divides into two daughter cells, is fundamental to all living organisms. Cytokinesis, the final step in cell division, begins with the formation of an actomyosin contractile ring, positioned midway between the segregated chromosomes. Constriction of the ring with concomitant membrane deposition in a spatiotemporal manner generates a cleavage furrow that physically separates the cytoplasm. Unique lipids with specific biophysical properties have been shown to localize to intercellular bridges (also called midbody) connecting the two dividing cells; however, their biological roles and delivery mechanisms remain largely unknown. In this study, we show that Ceramide phosphoethanolamine (CPE), the structural analog of sphingomyelin, has unique acyl chain anchors in spermatocytes and is essential for meiotic cytokinesis. The head group of CPE is also important for spermatogenesis. We find that aberrant central spindle and contractile ring behavior but not mislocalization of phosphatidylinositol phosphates (PIPs) at the plasma membrane is responsible for the male meiotic cytokinesis defect in CPE deficient animals. Further, we demonstrate the enrichment of CPE in multivesicular bodies marked by Rab7, which in turn localize to cleavage furrow. Volume electron microscopy analysis using correlative light and focused ion beam scanning electron microscopy shows that CPE enriched Rab7 positive endosomes are juxtaposed on contractile ring material. Correlative light and transmission electron microscopy reveal Rab7 positive endosomes as a multivesicular body-like organelle that releases its intraluminal vesicles in the vicinity of ingressing furrows. Genetic ablation of Rab7 or expression of dominant negative Rab11 results in significant meiotic cytokinesis defects. Our results imply that endosomal delivery of CPE to ingressing membranes is crucial for meiotic cytokinesis.

## Introduction

Eukaryotic cells display high diversity in lipid species that are chemically and structurally distinct. They are classified into eight major categories, each of which is further subdivided into classes and subclasses [1]. Lipids usually comprise of a polar head group that is linked by a structural backbone to hydrophobic tail groups composed of acyl chains. Lipid diversity that exists in cells arises from the large number of combinatorial chemical possibilities that the head and tail moieties can generate. Diversification of lipids to cope with evolving cellular complexity has resulted in thousands of lipid species with purported unique functions, most of which still remain unknown [2].

The head groups of each lipid class have been widely recognized for their biological function. For instance, the inositol head group of phosphatidylinositol phosphates (PIPs) and its modifications play critical roles in cellular signaling [3]. However, even within each lipid class, there are a number of molecularly distinct species that have identical head group but differ extensively in their acyl side chains, such as number of carbon atoms, number of double bonds, and the nature of the chemical linkage (e.g., ester or ether). An increasing number of studies are beginning to reveal the importance of acyl side chains by demonstrating that they selectively bind to target proteins and elicit biological function [4, 5]. For instance, a recent study showed that the acyl side chains of diacylglycerol significantly influence the recruitment of protein kinase C to the plasma membrane [6]. Similarly, ceramide chain length was shown to be critical for selective protein cargo sorting at endoplasmic reticulum (ER) exit sites in yeast [7]. The COPI machinery protein p24 bound specifically to N-stearoyl sphingomyelin (SM C18) and functioned as a cofactor to regulate COPI-dependent transport in the early secretory pathway. Further, this interaction depended on both the headgroup and the backbone of the sphingolipid [8].

Cytokinesis is the final step in cell division that divides one cell into two daughter cells. During cytokinesis, an actomyosin contractile ring is formed that is positioned midway between the segregated chromosomes. Narrowing of this ring with the addition of membrane in a spatiotemporally defined mode produces a cleavage furrow that physically divides the cytoplasm [9]. During somatic cytokinesis, the two daughter cells are interconnected *via* intercellular bridges prior to abscission. Although these structures are transient, they accumulate defined lipid species to mediate correct cell division [10–12]. Unlike somatic cytokinesis, in testis, the developing spermatogonial cells divide synchronously with an incomplete cytokinesis where all the daughter cells remain interconnected by cytoplasmic bridges. Thus, the spermatogonial cells develop as a syncytium and become separated from each other only at the end of spermatogenesis during spermatid individualization. The high degree of membrane curvature necessary for cleavage furrow ingression and stabilization of intercellular bridges (ICB)/cytoplasmic bridges necessitates lipid components with specific biophysical properties. While the presence of very long chain fatty acids (which are components of sphingolipids and glycosphingolipids) in cytokinesis has been shown, the relationship between membrane lipid composition, ring constriction and furrow ingression is still underexplored. Although sphingomyelins containing very long chain polyunsaturated fatty acids (VLC-PUFAs) were identified in various mammalian testes including humans, their direct role in spermatogenesis particularly in meiotic cytokinesis remains unknown [13, 14].

Ceramide phosphoethanolamine (CPE) is a structural analog of sphingomyelin (SM) in *Drosophila melanogaster*. CPE is synthesized in the luminal compartment of trans Golgi by ceramide phosphoethanolamine synthase (CPES) (Fig.1A). Previously, we have shown that *cpes* null mutants show significant late pupal lethality during development and only about 25% of mutants survive to adulthood. About 60% of *cpes* mutant adults have dorsal closure defects and all the mutant males are sterile. Aged *cpes* mutant adults display light inducible seizures and paralysis due to defective cortex glia [15]. In this study, we show that CPE has unique acyl chain anchors in spermatocytes and is essential for meiotic cytokinesis, spermatid polarity and individualization. The head group of CPE is also important for spermatogenesis. CPES expression in transit amplifying to the spermatocyte stage is essential for successful completion of spermatogenesis. We show that aberrant central spindle behavior, but not mislocalization of PIPs at the cleavage furrows is responsible for the meiotic cytokinesis defect in *cpes* mutants. Further, we show that endocytically retrieved CPE from plasma membrane is enriched in Rab7 positive multivesicular bodies that dock to the ingressing membranes and release intraluminal vesicles in their vicinity. Our results demonstrate the importance of CPE rich membrane addition at the cleavage furrow involving the endocytic pathway.

**Fig. 1.**
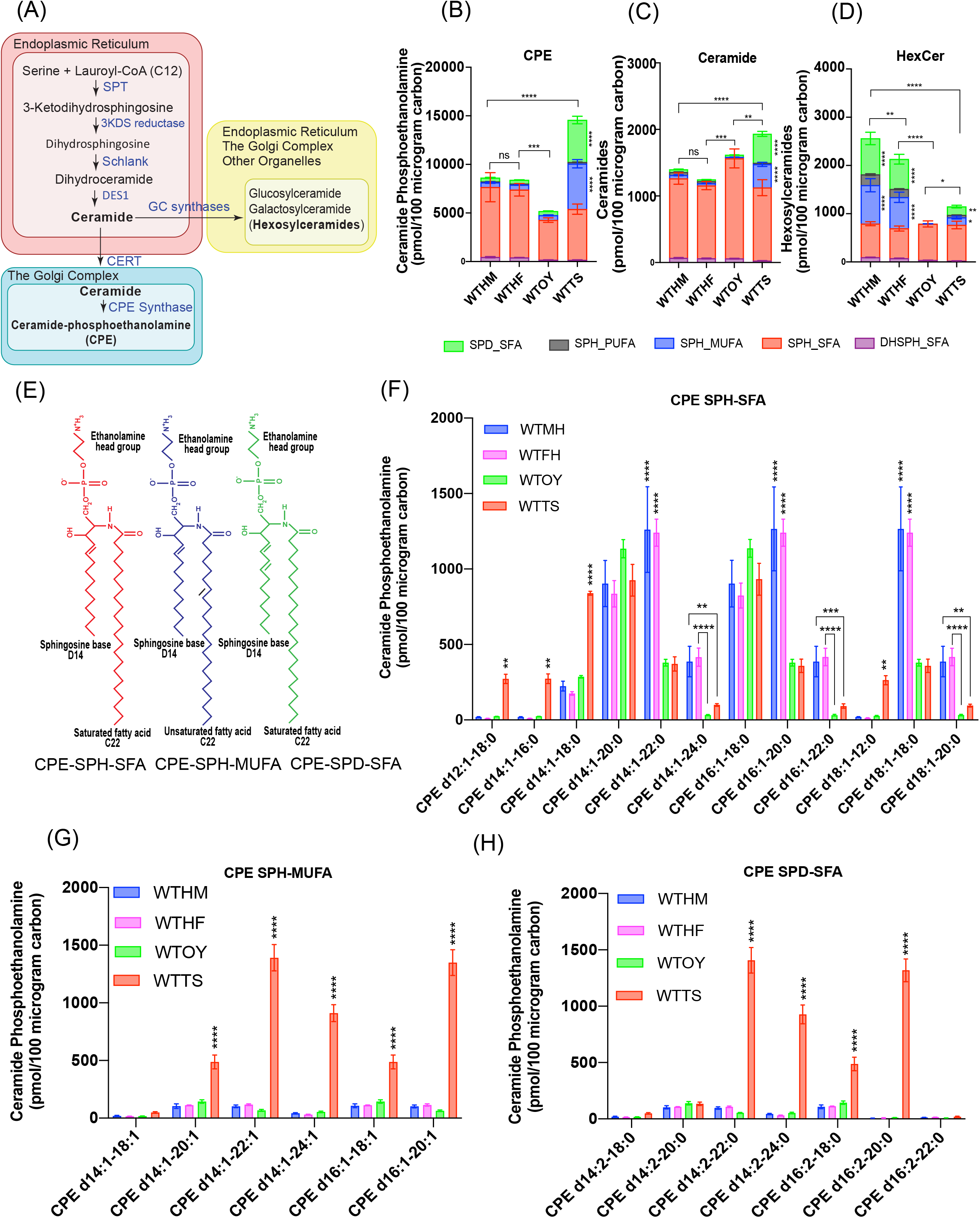
Sphingolipid analysis in wild type Drosophila tissues. (A) De novo sphingolipid biosynthetic pathway. De novo sphingolipid biosynthetic pathway begins with condensation of an amino acid serine and a fatty acid, lauryl-CoA into 3-ketodihydrosphinganine by serine palmitoyl transferase (SPT) complex in the membranes of endoplasmic reticulum (ER). 3-ketodihydrosphinganine (3KDS) is reduced to dihydrosphingosine by 3KDS reductase. Dihydrosphingosine is acylated to produce dihydroceramide by Schlank. Dihydroceramide is desaturated by Δ4 desaturase (DES1) resulting in the formation of ceramide. Ceramide acts as a precursor for the biosynthesis of complex sphingolipids both in the ER and Golgi complex. Ceramide is actively transported to Golgi by ceramide transfer protein (CERT). In the trans Golgi lumen, ceramide is converted to hexosylceramides by GC synthases and to CPE by CPE synthase. (B-D) Sphingolipids were extracted from wild type male and female heads and dissected ovaries and testes using Methanol:Chloroform method and subjected to SFC/MS/MS. Wild type heads from male flies (WTHM), wild type heads from female (WTHF), wild type ovary (WTOY) and wild type testis (WTTS). Sphingolipids were quantitated by measuring large fragments in MRM, and average amounts from 3 independent biological replicates are shown as picomoles (pmol) in 100 micrograms of carbon. Each sphingolipid species (CPE (1B), Ceramide (1C) & HexCer (1D)) was further subdivided into 5 major subspecies. Each of the subspecies differed in acyl chain length but similar in chemical properties: Dihydrosphingosine (DHSPH) base linked to saturated fatty acid (SFA) (DHSPH_SFA), Sphingosine (SPH) base linked to SFA (SPH_SFA), Sphingosine (SPH) base linked to Monounsaturated fatty acid (MUFA) (SPH_MUFA), Sphingosine (SPH) base linked to Polyunsaturated fatty acid (2-6 double bonds, PUFA) (SPH_PUFA) and Sphingadiene (SPD) with SFA (SPD_SFA). (E) Cartoon showing the chemical structure of three major CPE subspecies including CPE_SPH_SFA, CPE_SPH_MUFA, and CPE_SPD_SFA. (F) CPE subspecies within CPE_SPH_SFA that differed in number of carbon atoms in acyl chains of sphingosine base and fatty acid. (G) CPE subspecies within CPE_SPH_MUFA. (H) CPE subspecies within CPE_SPD_SFA. Statistical significance was calculated using mean, standard deviation (SD) and N in Prism 8. The 2way ANOVA multiple comparison was used to calculate P values where ****P ≤ 0.0001;***P ≤ 0.001; **P ≤ 0.01; *P ≤ 0.05 and ns P >0.05. Three independent biological replicates were performed for each sample and lipids were extracted from 400 heads or 250 pairs of ovaries or testes for each biological replicate.

## Results

### Sphingosine with VLC-mono-unsaturated fatty acid and sphingadiene with VLC-saturated fatty acid containing sphingolipids are enriched in the testis

To investigate sphingolipid species diversity and their role in *Drosophila* tissues, we performed supercritical fluid chromatography coupled to mass spectrometry (SFC/MS/MS) on lipids from wild type fly head, dissected ovary, and testis (Data S1). As shown in Fig.1B, CPE is a more abundant sphingolipid compared to ceramides (Fig.1C) and hexosylceramides (HexCer) (Fig.1D). The amount of CPE was significantly higher in testis compared to head and ovary (Fig.1B). In contrast hexosylceramides are significantly higher in heads compared to ovary and testis (Fig.1D).

Closer examination of acyl chain composition revealed that head and ovary derived CPEs are predominantly composed of sphingosine linked to saturated fatty acid (SPH_SFA) (Fig.1E and 1F). Interestingly, in addition to SPH_SFA, CPE in the testis is enriched in two distinct species with net two double bonds in their acyl chain composition including sphingosine (one double bond) linked to mono-unsaturated fatty acid (SPH_MUFA) (Fig.1E and 1G) and a sphingadiene (two double bonds) linked to saturated fatty acid (SPD_SFA) (Fig.1E and 1H). The fatty acid acyl chain lengths of these CPE species are predominantly composed of C20, C22, C24 and therefore could be classified as very long chain (VLC) sphingolipids. The sphingoid base acyl chain length is composed of d14 and d16 (Fig.1G and 1H). The corresponding ceramide precursors are also evident in the wild type testis samples indicating acyl chain diversity is likely introduced during ceramide synthesis but not after CPE synthesis (Fig.1C and Data S1). Remarkably, SPH_MUFA and SPD_SFA are also present in HexCer but at significantly lower levels in the testis compared to heads (Fig.1D vs Fig.1B). The fatty acid acyl chain length of HexCer_SPH_MUFA and HexCer_SPD_SFA showed that they are predominantly composed of C18 and C20 while the sphingoid base is composed of primarily d14 and d16 (Fig.S1B, S1D and S1E). HexCer with d14 sphingosine base and C22 PUFA (SPH-PUFA) is enriched in heads compared to testis (Fig.1D, Fig.S1C and S1E). Taken together, these results suggest that CPE in the testis and the HexCer in the central nervous system have unique acyl chain composition.

### CPE deficient males are sterile and show defects in male meiotic cytokinesis, spermatid polarity and individualization

To understand the biological significance of the unique CPE species in the testis, we reasoned that the *cpes* null mutant generated by us would serve as a suitable tool since *cpes* mutant males are sterile [15]. Immunostaining of *cpes* mutant testis with antibody to Vasa protein, a germ cell specific conserved RNA helicase [16], showed accumulation of germ cells at the tip of the testis (Fig. 2B) and DNA specific staining with DAPI showed absence of mature sperm in seminal vesicles (data not shown). To investigate cytological defects in *cpes* mutant testis, we performed live testis squash preparation followed by phase contrast microscopy. This method allows for easy identification of almost all of the major stages of spermatogenesis [17]. During meiotic divisions, chromosomes and mitochondria are equally partitioned to each of the four daughter cells. Immediately following meiosis, in the round spermatid stage, mitochondria in each daughter cell aggregate to form a large phase dark structure known as the nebenkern. Round spermatids (onion stage spermatids) are characterized by phase dark nebenkern and phase light nucleus at roughly equal size and at a ratio of 1:1 per daughter cell. The nuclear size of round spermatids are directly proportional to the chromatin content, hence any errors in chromosome segregation during meiosis can lead to variability in nuclear size [18, 19]. Similarly, defects in meiotic cytokinesis following normal chromosome segregation results in cells with two or four nuclei and an abnormally large nebenkern. This change can be easily visualized by phase contrast microscopy and is often used as a diagnostic to detect male meiotic defects [19–25]. As shown in Fig.2D, wild type round spermatids contained an equal sized phase light nucleus and a phase dark nebenkern at a ratio of 1:1 per cell within the cyst of 64 cells. In contrast, *cpes* mutant round spermatids showed four regularly sized nuclei (tetranucleate) and one abnormally large nebenkern, indicating a failure in cytokinesis in both meiosis I and II divisions (Fig.2E). Quantification of this defect suggest 80% of the round spermatids show tetranucleate and about 20% show two nuclei (Fig.2M). We did not observe single nucleated round spermatids *in* cpes mutant testes (Fig.2M). This defect is unique to male meiotic cytokinesis since larval neuroblasts undergoing asymmetrical mitotic division do not show defects in cytokinesis (Movie S1). Females are largely fertile indicating normal oogenesis (data not shown). Also, even during spermatogenesis we do not observe any defect in the formation of 16 cell staged spermatogonia/spermatocytes indicating that mitotic cytokinesis even in the testis of *cpes* mutants, is not compromised.

**Fig. 2.**
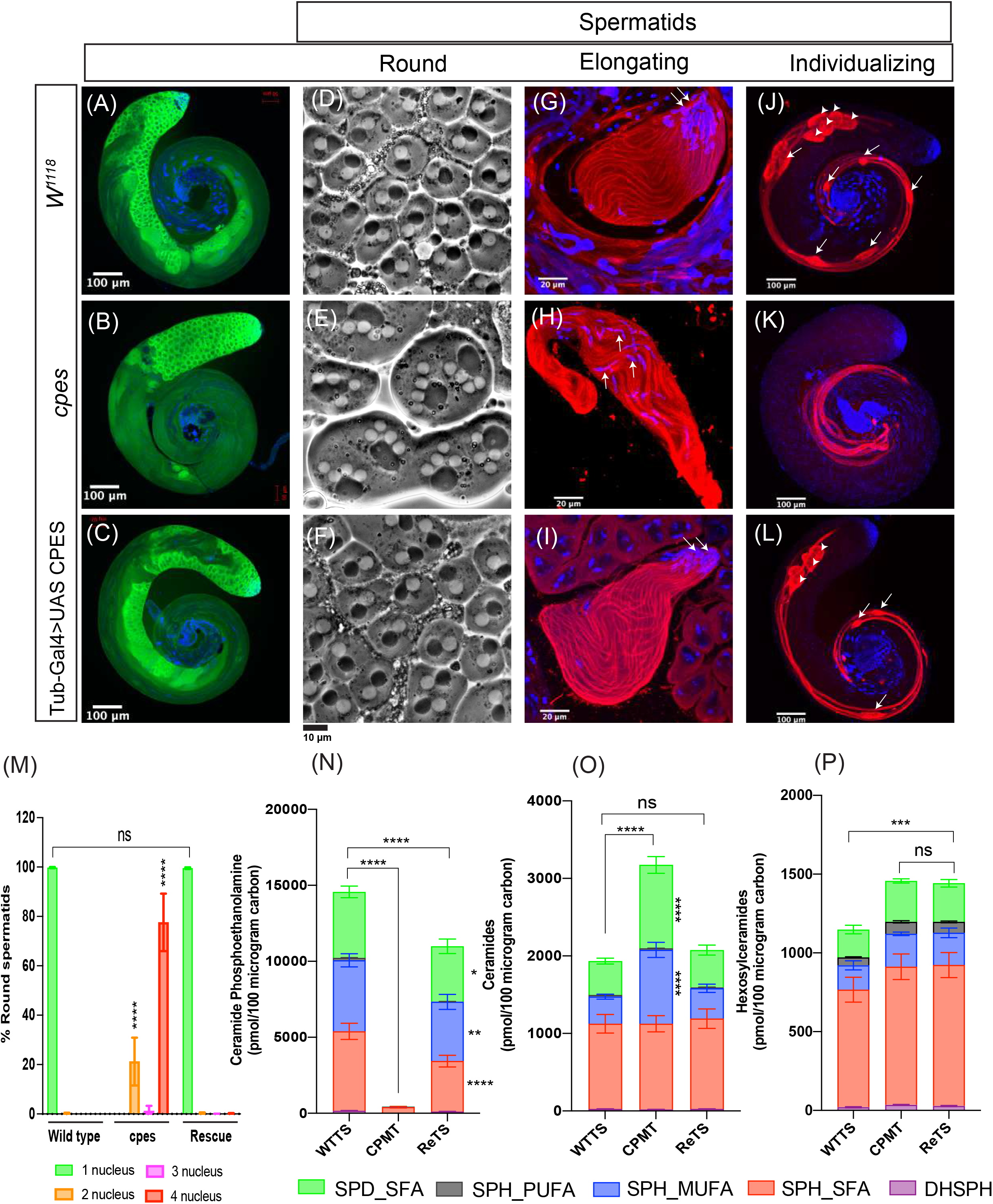
Cytological, immunofluorescence and sphingolipid analyses in cpes mutants. (A-C) Whole mount of Drosophila testis immunostained for germ cell specific Vasa protein primary antibody and Alexa Fluor 488 secondary antibody (Green) and DNA with DAPI (Blue) to visualize various stages of germ cells including stem cells, spermatogonia and spermatocytes (A) w^1118^, (B) cpes mutant and (C) Tub-Gal4>UAS-CPES rescue. (D-F) Phase contrast microscopy of live testis squash preparations to visualize round/onion stage spermatids (D) w^1118^, (E) cpes mutant and (F) Tub-Gal4>UAS-CPES rescue. (G-I) Testis squash preparation followed by fixation and immunostaining to visualize early elongating spermatids. Arrow indicates spermatid head stained with DAPI (Blue) for DNA. The beta tubulin primary antibody and Alexa Fluor 568 conjugated secondary antibody (Red) were used for immunostaining spermatid tails (axonemes), w^1118^ (G), cpes mutant (H) and Tub-Gal4>UAS-CPES rescue (I). (J-L) Whole mount Drosophila testis immunostained with cleaved caspase specific DCP-1 primary antibody and Alexa Fluor 568 secondary antibody (Red) to visualize elongated and individualizing spermatids. DAPI stains the DNA (Blue). Cystic bulges are shown with arrows and waste bags with arrow heads (J) w^1118^ (K) cpes mutant and (L) Tub-Gal4>UAS-CPES rescue. (M) Quantification of meiotic cytokinesis defects in cpes mutants. Round spermatid count from 3 independent experiments for W^1118^ (n=1600), cpes (n=300) and from 4 independent experiments for tubulin-Gal4 >UAS-CPES rescue (n=2000). (N-P) Sphingolipid analysis in lipid extracts of dissected testis, wild type testis (WTTS); cpes mutant testis (CPMT) and bam-Gal4>UAS-CPES Rescue testis (ReTS). (N) Comparison of CPE and its subspecies between w^1118^, cpes mutant and ReTS. (O) Comparison of ceramide and its subspecies between w^1118^, cpes mutant and ReTS. (P) Comparison of hexosylceramide subspecies between w^1118^, cpes mutant and ReTS. Statistical significance was calculated using mean, standard deviation (SD) and N in Prism 8. The 2way ANOVA multiple comparison was used to calculate P values where ****P ≤ 0.0001; ***P ≤ 0.001; **P ≤ 0.01; *P ≤ 0.05 and ns P >0.05. Three independent biological replicates were taken for each sample and 250 pairs of testes were used for lipid extraction from each biological replicate.

Following the round/onion stage, the 64 interconnected haploid cells in the cyst become highly polarized with the nuclei localizing to one end and the sperm tails (axonemes) growing in the opposite side. These elongating spermatid nuclei face towards the seminal vesicles and the tails grow towards the germline stem cell hub/tip of the testis. As shown in Fig.2G, wild type early elongating spermatids had all the nuclei facing one side and their axonemes growing to the opposite side indicating that they are highly polarized. In contrast, all the elongating spermatids in *cpes* mutant cysts appeared as sickle cell shaped with the nuclei present in the middle-bulged area (occasionally throughout the cyst) and their tails growing on both sides of the cyst (Fig.2H). This data indicates that spermatid polarity is compromised in *cpes* mutants.

Individualization is the final stage of spermatid differentiation wherein structures called the individualization complex (IC) containing actin cones assemble at the head/rostral end of elongated spermatids and start moving away from the nuclei towards the tail/caudal end of the cyst. As the IC travels it removes the cytoplasmic contents of the cyst and individualizes each spermatozoon with its own plasma membrane. The extruded cytoplasmic content is collected into a sac like structure called the ‘cystic bulge’ that forms around the IC. As this process proceeds and the IC and cystic bulge reach the end of the flagella, the actin cones and cytoplasmic contents are extruded in a waste bag, the contents of which are degraded later [26]. Activated caspases have a non-apoptotic role in cystic bulge and the waste bag of individualizing spermatids and are essential for successful spermatid individualization [27]. Immunostaining of wild type testis with cleaved *Drosophila* Caspase 1 (DCP-1) antibody showed elongated spermatids undergoing individualization that contained cystic bulges and waste bags (Fig.2J). In contrast, *cpes* mutant testis showed defectively elongated spermatids and was devoid of cystic bulges and waste bags indicating failure in spermatid individualization (Fig.2K).

The meiotic cytokinesis, spermatid polarity and individualization defects of *cpes* mutants are fully penetrant (100%) and ubiquitous expression of wild type cDNA copy of CPES completely rescued all male sterility phenotypes (Fig. 2C, F, I, L, M). As shown in Fig.2F, phase contrast imaging of live testis squash preparations showed 1:1 nebenkern to nucleus ratio indicating rescue of meiotic cytokinesis defect. Further, immunostaining of testis squash preparations with tubulin antibody and DAPI showed that spermatid polarity was completely restored wherein all the nuclei faced one side and their tails faced the other side of the cyst (Fig.2I). DCP-1 staining showed individualizing spermatids with cystic bulges and waste bags indicating normal individualization (Fig.2L).

To investigate the sphingolipid content in *cpes* mutant, we dissected wild type, *cpes* mutant and germ cell specific rescue (*bam*-Gal4> UAS-CPES) testes, sphingolipids were extracted and analyzed by SFC/MS/MS as described in the methods (Data S1). The mutant and rescue samples were processed at the same time as the wild type for mass spectrometry, hence the wild type testis sample is the same as in Fig.1 (Data S1). There was complete loss of CPE in *cpes* mutant testis (Fig.2N) and expression of CPES only in the germ cells was sufficient to restore CPE to near wild type levels. Loss of CPE synthesis in *cpes* mutants caused concurrent accumulation of ceramides particularly SPH_MUFA and SPD_SFA species (Fig.2O) and expression of CPES in germ cells reduced ceramide levels. HexCer levels were also slightly increased in *cpes* mutant testis (Fig.2P). Interestingly, expression of CPES in germ cells did not reduce the HexCer levels indicating their levels did not correlate with the *cpes* mutant phenotypes in the testis.

### CPES enzymatic activity is required for spermatogenesis and accumulation of ceramide is not the cause for spermatogenesis defects

To determine whether enzymatic activity of CPES is required for spermatogenesis we generated an active site mutant of CPES. CPES has a conserved amino acid motif D(X)_2_DG(X)_2_ (A/Y)R(X)_8-16_G(X)_3_D(X)_3_D in the CDP-alcohol phosphotransferase domain (CAPT) [28]. The final aspartates of this motif were shown to be essential for catalysis of human choline/ethanolamine phosphotransferase CEPT1 [29, 30]. We substituted the last two active site aspartates of CPES with alanine residues and subcloned into pUAST vector for generating transgenic flies. The UAS_CPES (DX_3_D to AX_3_A) mutant transgene was expressed in *cpes* mutant background using *bam*-Gal4 to test if rescue of spermatogenesis phenotypes could be rescued. As shown in Fig.S2A and B, the active site mutant did not rescue spermatogenesis phenotypes suggesting a crucial role for CPES enzymatic activity in spermatogenesis.

We next investigated whether accumulation of ceramide at the Golgi is responsible for the observed phenotypes. Ceramide is generated in the membranes of endoplasmic reticulum (ER) *via* the *de novo* biosynthetic pathway catalyzed by the rate limiting enzyme serine palmitoyltransferase (SPT) (Fig.1A). Subsequently ceramide is actively transported from ER to Golgi *via* ceramide transfer protein (DCERT) and thus absence of DCERT could prevent ceramide accumulation at the Golgi (Fig.1A) [31]. To investigate whether blocking ceramide transport from ER to Golgi could restore spermatogenesis, we generated *cpes* and *dcert^1^* double mutants and their testes were analyzed for meiotic cytokinesis and spermatid polarity phenotypes. However, as shown in Fig.S2C and D, *cpes*; *dcert^1^* double mutants did not rescue meiotic cytokinesis and spermatid polarity indicating ceramide accumulation at the Golgi (due to lack of CPES activity in the Golgi) may not be responsible for *cpes* mutant phenotypes. In *Drosophila*, transgenic expression of neutral ceramidase was shown to reduce ceramide levels *in vivo* [32]. We overexpressed ceramidase (UAS CDase) in germ cells using *bam*-Gal4 in *cpes* mutant background and testes from the resulting progeny were analyzed for the rescue of mutant phenotypes. However, expression of CDase did not rescue meiotic cytokinesis and spermatid polarity (Fig.S2E and F), suggesting that absence of CPE but not accumulation of ceramide is responsible for the phenotypes.

### Transcriptomic and genetic dissection of spatiotemporal role of CPES in spermatogenesis

Spermatogenesis is a highly conserved process across various taxa that is facilitated by highly dynamic transcriptome. An earlier study leveraged single cell RNAseq and unsupervised clustering to identify all the major cell classes of the sperm lineage and validated them with previously studied marker genes [33]. To characterize cell type specific transcriptional signatures in *cpes* mutants *in vivo*, we performed total RNA-seq with dissected testis from *w^1118^*, *cpes* mutant and *bam*-Gal4>UAS CPES rescue. Using Gene Set Enrichment analysis (GSEA) we compared our RNAseq results with the top 50 expressed genes from each cell type/cellular status that were previously classified to confirm the presence of germline stem cells (GSC), spermatogonia, spermatocytes, and spermatids (germ cells) as well as cyst stem cells, terminal epithelial cells and hub cells (somatic cells) [33]. We found that gene sets corresponding to GSC, early spermatogonia and late spermatogonial cells are enriched in *cpes* mutants compared to wild type and rescue testis (Fig.S3). In contrast, gene sets corresponding to early and later spermatocytes, early, and mature spermatids are enriched in wild type and rescue testis compared to *cpes* mutant testis (Fig.S3). These results suggested that CPES might play important roles during late spermatogonial, early and late spermatocyte stages for successful completion of spermatogenesis.

To determine the stage of germ cell differentiation at which CPES expression has an important role, we performed cell type/stage specific rescue experiments using UAS/Gal4 system in the *cpes* mutant background (Fig.3A). We first expressed UAS-CPES in cyst cells using C587-Gal4 to investigate a possible non-cell autonomous role. C587-Gal4 was expressed in all early somatic cells at the apical tip of the testis including somatic cyst stem cells and cyst cells (Fig.S4 D &H) [34]. As shown in Fig.3, expression of wild type CPES in cyst cells did not rescue meiotic cytokinesis, spermatid polarity or individualization defects (Fig.3 B, F and J respectively). We next expressed wild type UAS-CPES in germ cells using three different Gal4 drivers that have been previously shown to be specifically expressed in stem cells (*nos*-Gal4), transit amplifying spermatogonial cells (*bam*-Gal4) and early spermatocytes (*chif*-Gal4) (Fig.3A). Na*nos* (*nos*) gene encodes for an RNA-binding protein involved in formation of translational repressor complex. It is required for germ plasm organization, germline development, germline stem cell renewal and neuronal morphogenesis [35]. To specifically express a transgene of interest in male and female germline stem cells *nanos*-Gal4-VP16 was routinely used. This construct consists of 700bp *nos* promoter, Gal4-VP16 ORF, *nos* 3’ UTR and 500bp of 3’ genomic *nos* transcription unit. In male germ line *nos*-Gal4 primarily expresses in GSC and early spermatogonial stages [36]. Bag of marbles (*bam*) gene encodes for a fusome (a germ cell specific organelle) associated protein. The Bam protein is required for activation of the switch from spermatogonia to spermatocytes. The construct *bam*-Gal4-VP16 has been successfully used to drive the expression of transgenes in late spermatogonia and early spermatocytes in *Drosophila*. This construct consists of 900bp bam promoter, 500bp *bam* 5’UTR, Gal4:VP16 ORF and HSP70 3’UTR. Although transgenes driven with *bam*-Gal4-VP16 express in late spermatogonia through early spermatocytes, the peak expression was shown to occur during spermatocytes [37]. Gene trap Gal4 insertion collection screen identified *Chiffon*-Gal4 to drive transgene expression specifically in early spermatocytes and somatic cysts cells [38]. To verify the expression pattern of these Gal4 drivers in our experimental settings we have crossed *nos*-Gal4 (BDSC#4937) with various pUASP/pUAST-GFP tagged proteins including pUASP-*alpha-Tubulin*-GFP, pUAST-EGFP, and pUAST-PLC*δ*-PH-EGFP. As expected, *nos*-Gal4 expression is restricted to early germ cells present at the tip of the testis including germline stem cells (GSCs) and early spermatogonial cells (Fig.S4A, E & I). On the contrary, *bam*-Gal4 expression is strong just below the testis tip where spermatogonia and early spermatocytes are expected to be present (Fig.S4B, F & J). However, depending on the stability of the expressed protein *bam*-Gal4 mediated expression is either restricted to spermatogonia and early spermatocytes (eg. PLC*δ*-PH-EGFP, Fig.S4J) or persisted through later stages including early, later spermatocytes and even in spermatids (eg. *Alpha tubulin* and EGFP, Fig.S4B & F). Expression of *chif*-Gal4 expression was detectable in spermatocytes (Fig.S4C) but more strongly in elongated spermatids (Fig.S4G & K). Interestingly, As shown in Fig.3, we observed varying degree of rescue, depending on the germ cell differentiation stage at which UAS-CPES was expressed. The *nos*-Gal4 dependent expression of UAS-CPES partially rescued both meiotic cytokinesis (Fig.3C and N) and spermatid polarity (Fig.3G) however, individualizing spermatids lacked cystic bulges and waste bags indicating individualization was not fully rescued (Fig.3K). The *bam*-Gal4 dependent expression of UAS-CPES in transit-amplifying cells completely rescued meiotic cytokinesis (Fig.3D, and N) and spermatid polarity, and individualizing spermatids contained cystic bulges and waste bags (Fig.3H and L). In contrast, only partial rescue of meiotic cytokinesis was observed when UAS-CPES was expressed in spermatocytes using *chif*-Gal4 (Fig.3E & 3N). Although spermatid polarity was completely rescued, the individualizing spermatids largely lacked cystic bulges and waste bags indicating that individualization was defective (Fig.3I and M).

**Fig. 3.**
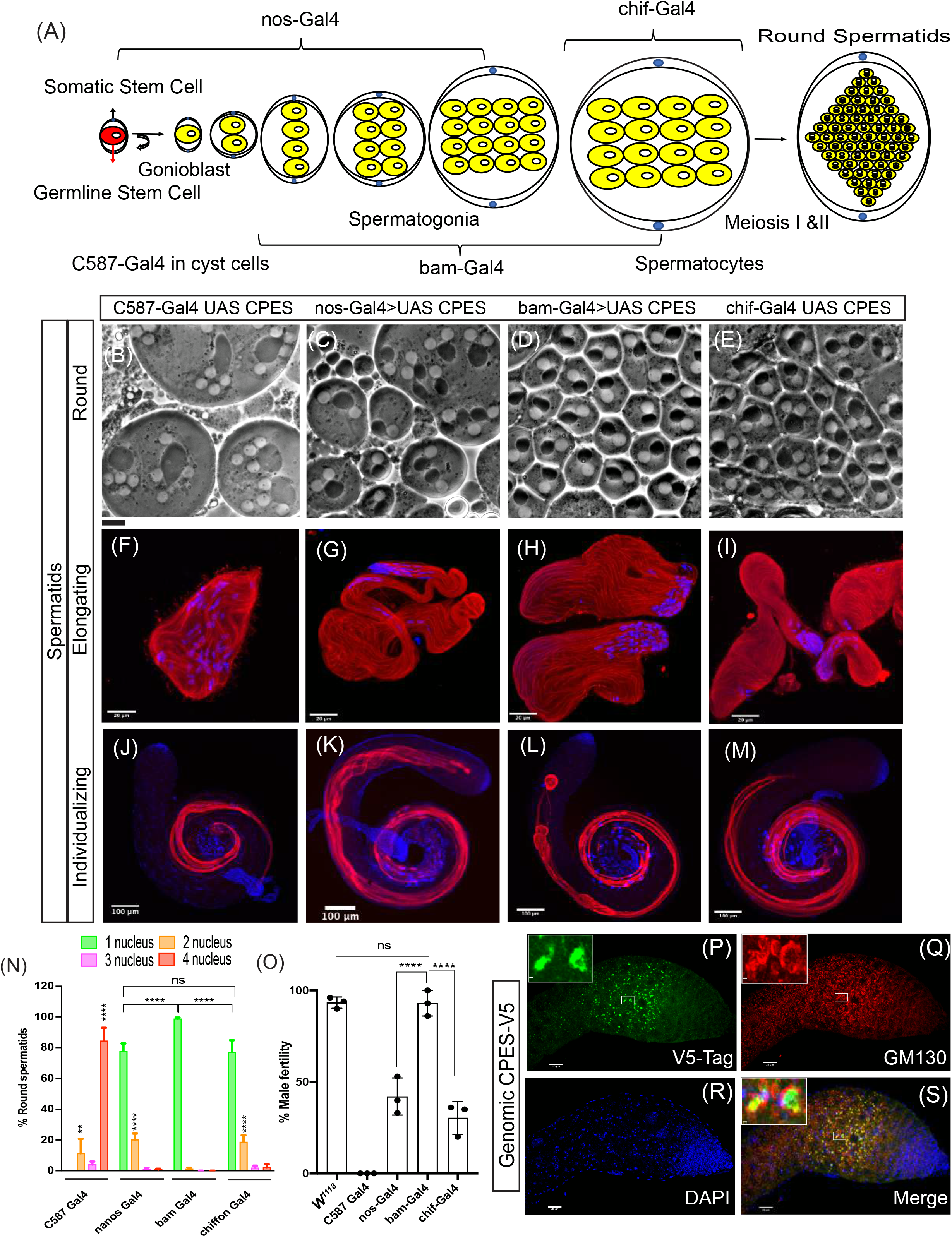
Genetic dissection of spatiotemporal role of CPES in spermatogenesis. (A) Schematic illustration of early germ cell differentiation and stage specific Gal4 expression patterns. (B-E) Phase contrast microscopy of round spermatids in live testis squash preparations. (F-I) Early elongating spermatids immunostained for beta tubulin primary antibody (Red) and DAPI for DNA (blue) (J-M) Individualizing spermatids immunostained for cleaved caspase DCP-1 antibody (Red) and DAPI for DNA(Blue). (B, F and J) C587-Gal4 drives the expression of UAS-CPES in cyst cells of cpes mutants. (C, G and K) Nanos-Gal4 drives the expression of UAS-CPES in GSCs and spermatogonia. (D, H & L) bam-Gal4 drives the expression of UAS-CPES in 4-16 cell stages of spermatogonia and spermatocytes (E, I & M) chif-Gal4 drives the expression of UAS-CPES in early spermatocytes. (N) Measurement of cytokinetic defects in testis squash preparations in cyst and germ cell specific rescue samples. Round spermatid count from 4 independent experiments for C587-Gal4 >UAS-CPES (n=500), 6 independent experiments nos-Gal4 >UAS-CPES (n=2500), 5 independent experiments for bam-Gal4 >UAS-CPES (n=2500), and 6 independent experiments for chif-Gal4 >UAS-CPES (n=3000). (O) Male fertility test where individual male from each of the UAS-CPES rescue experiment was crossed with 3 wild type females. The vials which produced at least 5 pupae or progeny were counted as fertile and plotted as percent fertility compared to wild type males. (P-S) Whole mount immunostaining followed by confocal imaging of genomic CPES flies where CPES is tagged with V5 at the C-terminus (red). GM130 is a cis-Golgi marker (green) and DAPI (blue) stains the DNA.

To determine the extent to which male fertility was rescued by cell type specific CPES expression compared to wild type, we performed male fertility tests. As shown in Fig.3O, expression of UAS CPE in transit amplifying spermatogonial cells and spermatocytes (*bam*-Gal4) rescued male fertility to wild type levels. In contrast only partial rescue was observed when UAS-CPES was expressed only in stem cells (*nos*-Gal4) or in spermatocytes (*chif*-Gal4). Expression of UAS-CPES in cyst cells (C587-Gal4) did not rescue male sterility as all males were sterile (Fig.3O). To determine *in viv*o CPES localization more precisely at the protein level, we have generated a construct with a 6437 bp extended CPES genomic fragment that was modified by recombineering to carry V5-tag at the C-terminus [15]. Transgenic flies expressing genomic CPES tagged with V5 at the C-terminus fully rescued lethality, photosensitive epilepsy and all other phenotypes including spermatogenesis defects. Immunostaining followed by confocal imaging showed that CPES_V5 was highly expressed in the cells that are present just below the tip of the testis (Fig.3P and S). This area of the testis is known to be enriched in late spermatogonia and early spermatocytes [17].

Taken together these results show that testis specific CPE generation by CPES from late spermatogonia stage to spermatocyte stage is crucial for successful meiotic cytokinesis, spermatid polarity and spermatid individualization.

### Insect CPE Synthases but not Sphingomyelin Synthase rescue male sterility defects

Previously we have shown that CPE has an important structural role in cortex glial plasma membranes [15]. The cortex glial plasma membranes are severely defective in *cpes* mutants leading to loss of cortex glia neuronal cell body interactions during development and increased susceptibility to light inducible epilepsy in adults. Further, we have shown that overexpression of human sphingomyelin synthase 1 (hSMS1) that produces SM instead of CPE was sufficient to rescue cortex glia plasma membrane defects via establishment of detergent resistant membranes. We wondered whether overexpression of hSMS1 could also rescue the spermatogenesis phenotypes of *cpes* mutant. To this end, UAS-hSMS1 was ubiquitously expressed using tubulin-Gal4 in *cpes*, the adult male testes were dissected, and live testis squash preparations were observed using phase contrast microscopy. However, as shown in Fig.S5A, hSMS1 overexpression did not rescue male meiotic cytokinesis. Further, immunostaining analysis showed that spermatid polarity was also not rescued (Fig.S5E) suggesting an important role for the CPE head group in spermatogenesis. We next investigated whether expression of other insect derived CPES could rescue *cpes* mutant phenotypes. To test this, we ubiquitously expressed *Aedes aegypti* (Yellow fever mosquito, amino acid sequence shows 60% identity and 77% similarity), and *Bombyx mori* (domestic silk moth, amino acid sequence shows 43% identity and 58% similarity) derived UAS-CPES homologs in *cpes* mutant background using tubulin-Gal4. As shown in Fig.S5 mosquito and silk moth CPES were able to completely rescue defects in meiotic cytokinesis (Fig.S5B-D) and spermatid polarity (Fig.S5F and G respectively).

To investigate the nature of sphingolipid species synthesized by hSMS1, *Aedes aegypti*, *Bombyx mori* derived CPES, we specifically expressed these transgenes in germ cells using *vasa*-Gal4, their male reproductive system was dissected, lipids were extracted and subjected to sphingolipid analysis using SFC/MS/MS as described in the methods section (Data S2). As shown in Fig.S5H germ cell specific expression of *Drosophila* CPES, *Bombyx mori* CPES completely restored CPE to the wild type amounts. *Aedes aegypti* CPES also synthesized significant amount of CPE but to a lower amount than *Drosophila* CPES. However, all three distinct CPE species are synthesized by *Aedes aegypti* CPES and *Bombyx* CPES (Fig.S5H). As anticipated, hSMS1 did not synthesize CPE, but produced significant amount of SM (Fig.S5K). However, the amount of SM is relatively low compared to CPE, perhaps due to limited expression/stability of hSMS1 in germ cells (vasa-Gal4). Indeed, ubiquitous expression of hSMS1 using actin-Gal4 driver significantly increases the SM amounts (Fig.S5L). However, expression of hSMS1 with ubiquitous drivers like actin-Gal4 and tubulin-Gal4 did not rescue the spermatogenesis (Fig.S5A and E; data not shown for actin-Gal4) suggesting the importance of the CPE head group. We next analyzed the ceramide content in these samples and found that ceramide levels were significantly high in *cpes* mutants and hSMS1 rescue compared to *Drosophila* CPES rescue (Fig.S5I). However, as discussed in Fig.S2 higher ceramide content might not contribute to the cytokinetic defects in *cpes* mutants. Similarly, higher amounts of HexCer in *Aedes aegypti* CPES rescue and *Bombyx mori* CPES rescue also did not correlate with the rescue of spermatogenesis phenotypes (Fig.S5J). Taken together these results suggest that the *Drosophila* spermatogenesis is strictly dependent on CPE and the head group of CPE play an important role.

### Aberrant central spindle behavior but not localization of PIPs at the plasma membrane is responsible for defective meiotic cytokinesis in *cpes* mutants

Precise organization of central spindle microtubule is not only required for initial cleavage furrow formation but also for maintenance of contractile structures during furrow ingression. Several microtubule interacting proteins including Fascetto, kinesin 6 family member MKLP1/Pavarotti (pav), chromokinesin klp3A, Orbit etc., are enriched in the central spindle midzone and are essential for cytokinesis [39]. To determine the transcriptional signatures relating to spindle organization and function in our RNAseq data, we performed GSEA pathway comparisons between *w^1118^* (WT), *cpes* (KO) and rescue (RES) samples. As shown in Fig.S6A. we found several pathways including spindle organization, metabolism, endocytosis, signaling, male meiotic cytokinesis, spermatid differentiation, sperm individualization etc., are significantly altered in *cpes* mutants (Fig. S6A). Red bars in the Fig. S6A indicate gene sets enriched in *cpes* mutants and blue bars indicate gene sets enriched WT & RES. GSEA showed many genes involved in spindle elongation and spindle organization are positively correlated with the mutant phenotype (Fig. S6B & E). The heat map comparison for genes involved in spindle organization and elongation are shown (Fig.S6). Several of the genes implicated in spindle organization have also been annotated under those for spindle elongation and hence appear under both categories. However, as shown in Fig.S3 relative enrichment of earlier stages compared to later stages limits the accurate prediction of altered pathways in *cpes* mutants.

To better study spindle behavior during male meiotic cytokinesis *in vivo*, we performed live imaging on isolated cysts. We first dissected the testis in M3 insect cell culture media and cut open the muscle sheath to release intact spermatocyte cysts into the media. Subsequently, we transferred spermatocyte cysts to poly-D-lysine coated cover glass dishes and performed live imaging using Andor spinning disk confocal microscopy as described in the methods section. We chose mCherry tagged microtubule cross linking protein Feo (Ubi-p63E-*feo*-mCherry) as a marker for central spindle. Feo-mCherry was shown to accumulate at the anaphase B and telophase central spindle [40]. We also used spaghetti squash (sqh) gene encoding myosin II regulatory light chain (RLC) tagged to GFP (*sqh*-GFP-RLC) as a marker for the actin-myosin based structure known as contractile ring that assembles during late anaphase [41]. The actin and myosin ring positioned midway between two spindle poles is thought to drive the formation of the cleavage furrow during telophase. As shown in Movie S2, Fig.4A, the actomyosin ring (*sqh*-GFP-RLC) quickly assembles around the central spindle during late anaphase and both simultaneously constrict through early to late telophase. At the end of late telophase *feo*-mCherry disassembles however, *sqh*-GFP-RLC remains and becomes part of ring canal/cytoplasmic bridges between daughter cells (Movie S2).

**Fig. 4.**
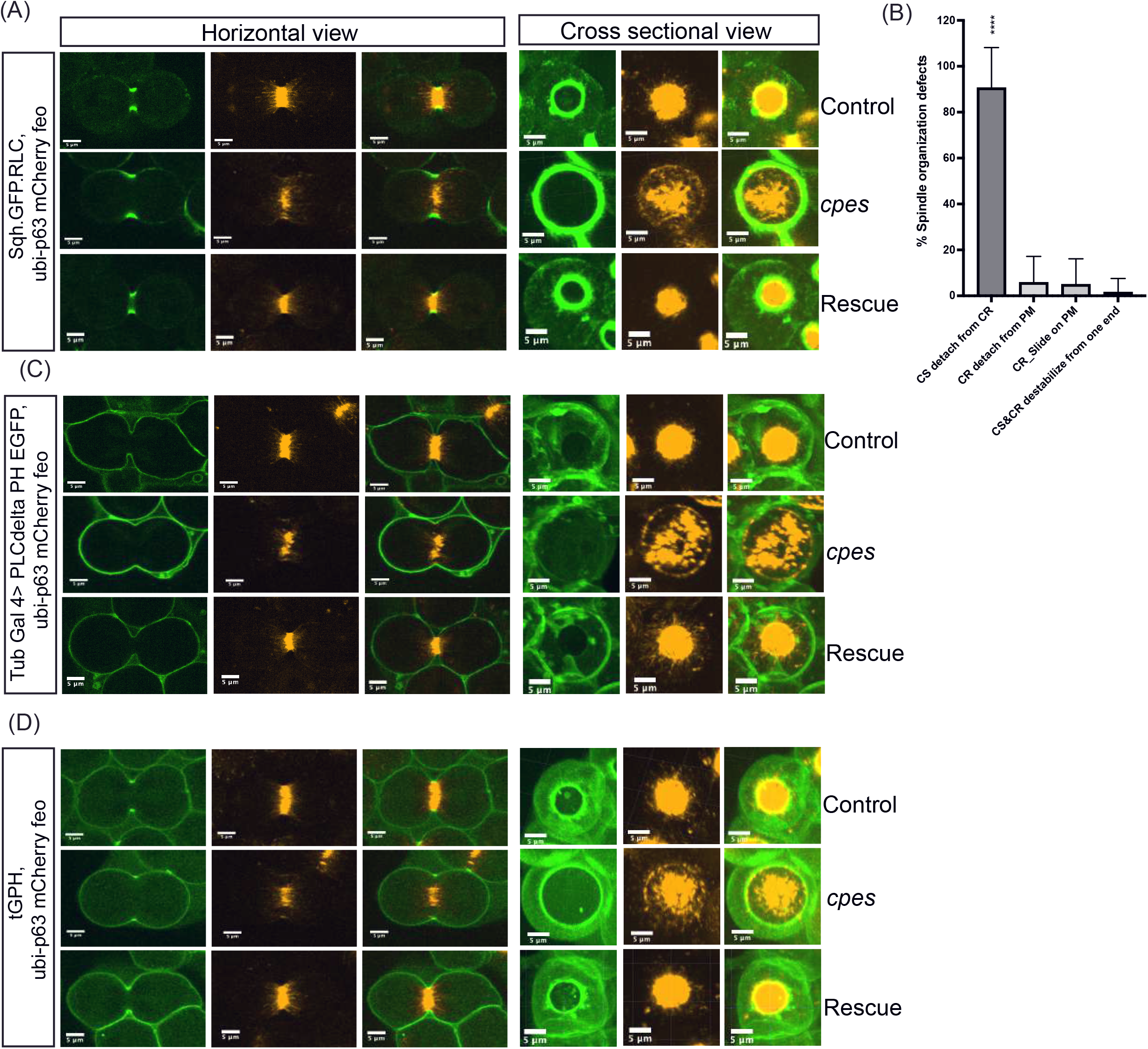
Aberrant central spindle behavior but not localization of PIPs at the plasma membrane is responsible for meiotic cytokinesis defect in cpes mutants. (A, C & D) Snapshots of live spermatocytes undergoing meiosis I cytokinesis. In horizontal view (left), one slice of a z-stack was shown. In cross-sectional view across cleavage furrow, maximum intensity projection image was shown (right). Control spermatocytes (top row), cpes mutant spermatocytes (middle row) and tubulin-Gal4>UAS-CPES rescue (bottom row). (A) Spermatocytes expressing myosin II regulatory light chain tagged to GFP under its own promoter (sqh) that localizes to contractile ring and mCherry tagged fascetto (feo) under the control of Ubi-p63E promoter that binds to central spindle microtubules. (B) Quantification of spindle organization defects, central spindle (CS) detachment from contractile ring (CR), detachment of CR from plasma membrane (PM), CR slides on PM and CS and CR destabilizing from end (C) Spermatocytes expressing GFP tagged pleckstrin homology (PH) domain of PLCD (UAS-PLCδ-PH-EGFP) under the control of Tub-Gal4 that localizes to plasma membrane in the presence of phosphatidylinositol-4,5-bisphosphate (PIP2) and Ubi-p63E-feo-mCherry. (D) Spermatocytes expressing GFP tagged PH domain of steppke/Grp1 under the control of alpha Tub84B promotor (tGPH) that localizes to plasma membrane in the presence of phosphatidylinositol-3,4,5-triphosphate (PIP3) and Ubi-p63E-Feo-mCherry.

Interestingly, in *cpes* mutants, the initial assembly of central spindle and actomyosin ring occurred normally (Movie S3) however, central spindle and actomyosin ring constriction became uneven since there is barely any furrow ingression and led to substantial disengagement between the contractile ring and the central spindle during meiosis (Fig.4A). As a result, the central spindle and actomyosin ring disassembled prematurely (Movie S3). Quantification of this behavior suggested that more than 80% of the spermatocytes showed detachment of the contractile ring and central spindle (Fig.4B). Other minor variations included the contractile ring detaching from the plasma membrane, contractile ring sliding and dumping to one end and the contractile ring and central spindle being destabilized simultaneously at one end (Fig.4B). Expression of UAS-CPE in germ cells was sufficient to rescue this phenotype (Movie S4, Fig.4A).

Previous studies have shown that phosphatidylinositols (PIP) play critical roles in somatic and meiotic cytokinesis [42]. Further, in mammalian (HeLa) cells undergoing mitosis, sphingomyelin-rich lipid domains in the outer leaflet of cleavage furrow were shown to be required for accumulation of PIP2 to the cleavage furrow which in turn is required for proper translocation of RhoA and progression of cytokinesis. We wondered whether the distribution of PIP at the plasma membrane or at the cleavage furrow is altered in *cpes* mutants during meiosis. To investigate PIP distribution in *cpes* mutant membranes we used PIP2 reporter UAS-PLC*δ*-PH-EGFP (UAS regulatory sequence drives expression of EGFP fused to the PH domain of human PLCdelta 1) and PIP3 reporter tGPH (an alphaTub84B promoter drives the expression of EGFP fused to the PH domain of Steppke) [43, 44]. The PIP2 reporter PLC*δ*-PH-EGFP was shown to uniformly localize to the entire membrane including cleavage furrow in *Drosophila* spermatocytes [45]. Indeed, we have seen similar uniform distribution in both control and *cpes* mutant spermatocyte plasma membranes indicating normal distribution of PIP2 (Movie S5 and S6, Fig.4C). Live imaging analysis on *cpes* mutant spermatocytes expressing PLC*δ*-PH-EGFP revealed normal cleavage furrow initiation, but progression of furrow failed due to the disconnect between central spindle and the furrow (Movie S6, Fig.4C). Expression of wild type CPES restored this phenotype (Movie S7, Fig.4C). Of note, PLC*δ*-PH-EGFP localization to spermatocyte plasma membrane was significantly reduced upon loss of cyst cells although this acute reduction did not prevent spermatocytes from completing cytokinesis during meiosis 1. We next investigated the PIP3 distribution in control meiotic spermatocytes using tGPH. Interestingly, we found that tGPH localized to plasma membranes and became significantly enriched at cleavage furrows (Movie S8, Fig.4D). Of note, even here, we have observed that plasma membrane localization and cleavage furrow enrichment of tGPH strictly depended on the intact cyst. While it is currently unknown if PIP3 localization to the spermatocyte cleavage furrows has any impact on cytokinesis, a recent study has shown that cytohesin Steppke reduces tissue tension by inhibiting the actomyosin activity at adherens junctions and actomyosin network assembly is necessary and sufficient for local Steppke accumulation in *Drosophila* embryos [46]. Next, we wondered if cleavage furrow enrichment of PIP3 is altered in *cpes* mutants. However, as shown in Movie S9, Fig.4D, PIP3 localization to the cleavage furrow was not altered but again there was a disconnect between cleavage furrow and central spindle during cleavage furrow ingression. Expression of wild type CPES restored this phenotype (Movie S10; Fig.4D). Overall, these observations suggest that aberrant spindle behavior but not the localization of PIPs to the plasma membrane or cleavage furrow, is responsible for meiotic cytokinesis defect in *cpes* mutants.

### CPE endocytosis and targeting to cleavage furrows occurs during meiotic cytokinesis

To determine how CPE is promoting the synergy between central spindle constriction and cleavage furrow ingression during cytokinesis we have utilized recombinant CPE binding protein tagged with mCherry as a reporter for visualization of CPE on spermatocyte plasma membranes. The mushroom-derived protein of the aegerolysin family pleurotolysin A2 (PlyA2) was shown to specifically bind to CPE and was demonstrated to be a versatile tool for visualizing CPE in live *Drosophila* tissues [47]. We have reevaluated the efficacy of recombinant PlyA2-mCherry protein in detecting endogenous CPE on live spermatocyte membranes. The wild type and *cpes* mutant spermatocytes were treated with 10 μg/ml of PlyA2-mCherry in M3 insect media for 1hour at room temperature, washed with fresh M3 media, and imaged on an Andor spinning disk confocal microscope. As shown in Fig.5A-D, PlyA2-mCherry specifically bound to the plasma membranes of wild type spermatocytes but not to the *cpes* mutant spermatocytes showing specificity of this protein binding to CPE. Interestingly, we found that PlyA2-mCherry was endocytosed in wild type spermatocytes (Fig.5A). To determine the nature of these endosomes where CPE is enriched, we performed live imaging on spermatocytes expressing endogenous EYFP-MYC tagged Rab proteins [48]. The small GTPases Rab4, Rab7 and Rab11 were shown to mark early, late and recycling endosomes respectively [49–51] while Rab6 was shown to be associated with the Golgi membranes [52]. The spermatocytes expressing individual EYFP-Rab proteins were treated with recombinant PlyA2-mCherry and imaged. As shown in Fig.5G, M and P, PlyA2-mCherry colocalized with Rab4, Rab7 and some of the Rab11 endosomes respectively. Colocalization coefficient measurements further support these observations (Fig.5Q). However, PlyA2-mCherry containing structures did not colocalize with Rab 6 positive Golgi membranes (Fig.5J). These results suggested that CPE on spermatocyte plasma membranes is actively targeted to the endocytic pathway. We next wondered whether endosomes carrying CPE migrate to cleavage furrows during meiotic cytokinesis. EYFP-Rab 7 and EYFP-Rab11 were strongly expressed in spermatocytes compared to Rab4 and therefore we followed their dynamics during meiotic cytokinesis. Strikingly, as shown in Fig.6A-F, & Movie.S11, PlyA2-mCherry containing Rab7 positive endosomes were actively targeted to cleavage furrows. Similar EYFP-Rab7 localization was seen in intact cysts (Fig.6G-I). The PlyA2-mCherry also colocalized to some of the EYFP-Rab11 recycling endosomes which in turn localized to ingressing membranes at the cleavage furrow (Fig.6J-O and Movie S12). Overall, the endocytosis of CPE and targeting to the ingressing cleavage furrow *via* Rab7 and Rab11 marked endosomes indicates their importance in delivery of lipids to the ingressing membranes of the furrow and coordination of this event with proper spindle behavior and meiotic cytokinesis.

**Fig. 5.**
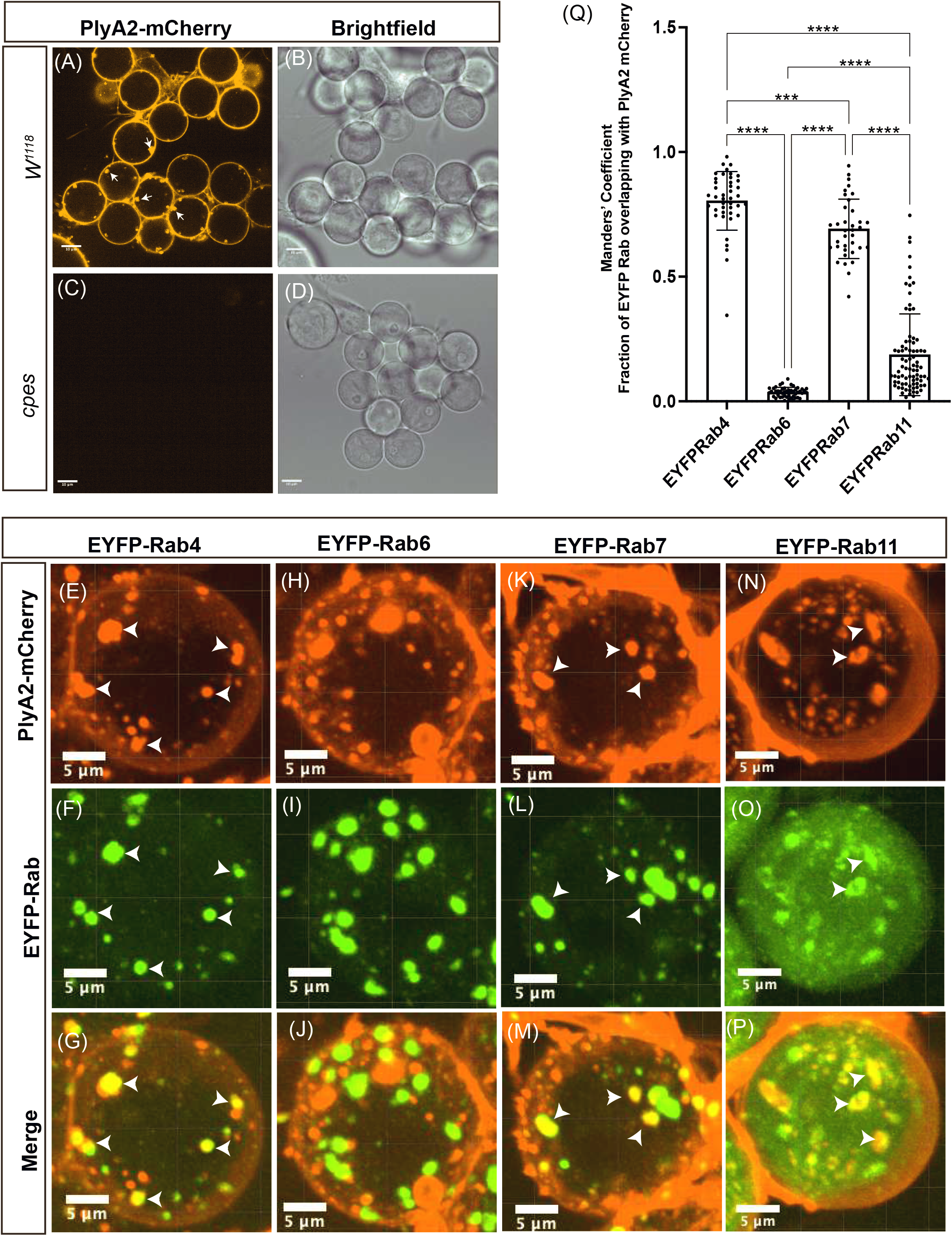
Ceramide phosphoethanolamine (CPE) localizes to plasma membrane and endosomes. (A-D) Live spermatocytes were incubated with purified PlyA2-mCherry (10 μg/ml) for 30 min followed by washes with M3 insect media (2x) and imaged using spinning disk confocal microscopy. (A and B) w^1118^ spermatocytes, (C and D) cpes spermatocytes. (E-P) Live spermatocytes expressing endogenous EYFP tagged Rab proteins were incubated with PlyA2-mCherry for one hour followed by washes with M3 insect media (2x) and imaged using spinning disk confocal microscopy. (E-G) EYFP-Rab4 spermatocytes, (H-J) EYFP-Rab6 spermatocytes, (K-M) EYFP-Rab7 spermatocytes, (N-P) EYFP-Rab11 spermatocytes. Arrowheads indicate colocalizing structures. (Q) Colocalization coefficient (Mander’s Coefficient) measurements showing fraction of EYFP Rab proteins colocalize with internalized PlyA2-mCherry. Mander’s coefficient was determined using imageJ plugin JACoP. Each dot in the graph represents a 2D image consisting of 8-16 spermatocytes, Rab4 (n=42), Rab6 (n=47), Rab7(n=36) and Rab11 (n=81). Statistical significance was calculated using mean, standard deviation (SD) and N in Prism 9. The ordinary one-way Anova multiple comparison was used to calculate P values where ****P ≤ 0.0001;***P ≤ 0.001; **P ≤ 0.01; *P ≤ 0.05 and ns P >0.05.

**Fig. 6.**
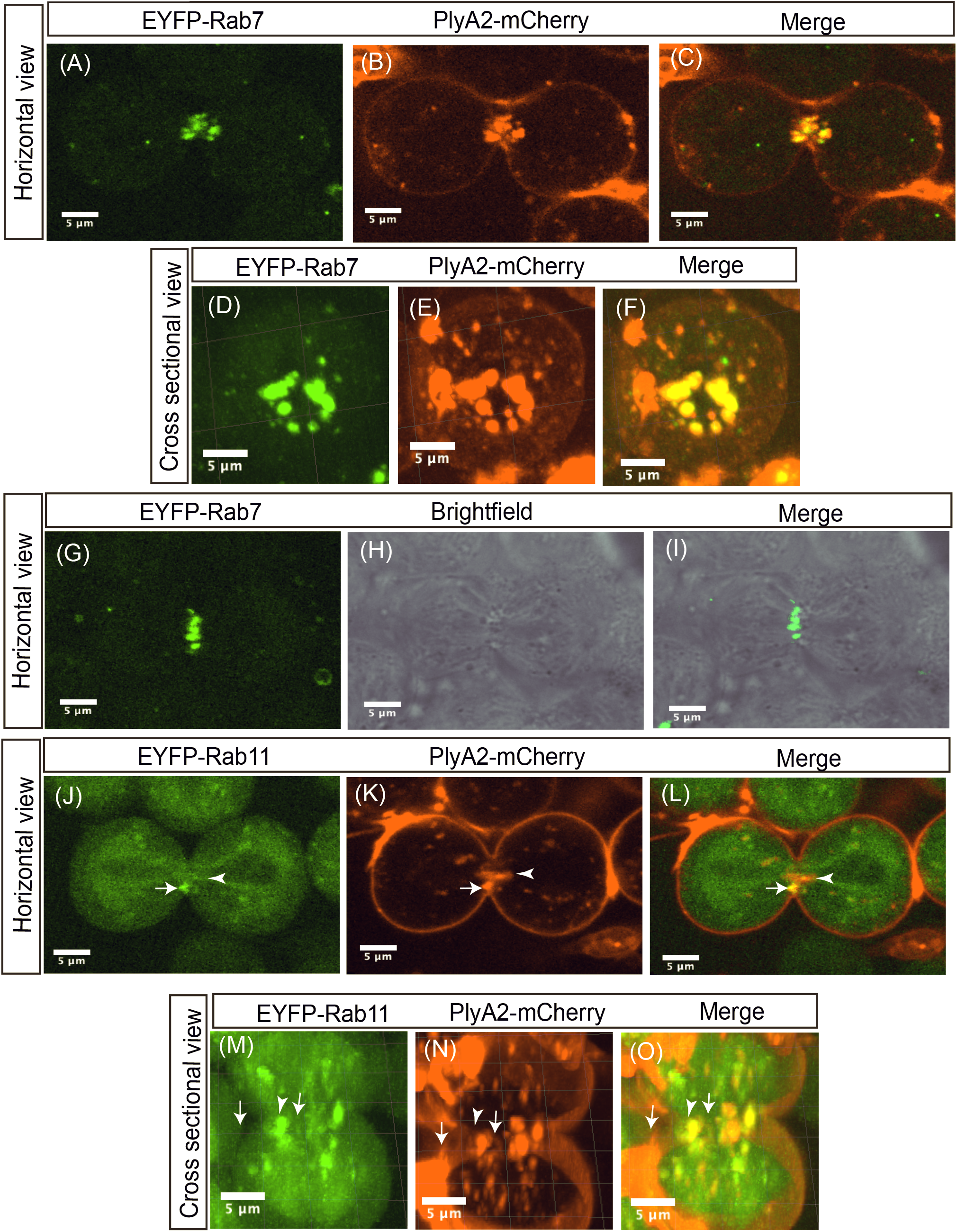
CPE containing endosomes translocate to cleavage furrows during male meiotic cytokinesis. (A-F) The EYFP-Rab7 expressing spermatocytes are treated with PlyA2-mCherry and snapshots of spermatocytes undergoing meiosis I cytokinesis are shown. (A-C) Horizontal view of one z slice is shown (D-F) Cross sectional view of a spermatocyte undergoing cytokinesis, maximum intensity projection image was shown. (G-I) Localization of EYFP-Rab7 in intact cysts where spermatocytes are encapsulated by cyst cells. Horizontal view of a single z slice is shown. (J-O) EYFP-Rab11 expressing spermatocytes were treated with PlyA2-mCherry and meiosis I cytokinesis was imaged (J-L) Horizontal view of single z slice is shown. (M-O) Cross sectional view, maximum intensity projection image is shown. White arrows indicate colocalization and white arrowheads indicate lack of colocalization.

To further verify the functional significance of endosome mediated CPE trafficking in cytokinesis we performed genetic experiments involving conditional expression of EYFP-Rab11DN (BDSC#) proteins and Rab7 mutants [53]. Rab11 null mutants die early during development [54]. In that study, it was demonstrated that when spermatocytes enter meiosis 1 as the Golgi disassembles, the Rab11 was shown to be associated with the ER compartment. Rab11 also localizes to vesicles at the cell poles during anaphase and early telophase and at the cell equator during mid- and late telophase. Larval testes of escaper flies carrying Rab11 semi lethal transheterozygote alleles showed abnormal constriction of the contractile rings and a failure of cytokinesis in 10-30% of the flies. We chose to conditionally express dominant negative EYFP-Rab11^S25N^ ubiquitously using Gal80^ts^ and tub-Gal4 to study cytokinetic defects in spermatocytes. The expression of EYFP-Rab11S25N (Rab11 DN) in late 3^rd^ instar larvae was induced by incubation at 30°C for 4 days. The late pupal gonads were dissected, and cytological analysis was performed to measure percent cytokinetic defects. Consistent with previously published results [54, 55], we found significant increase in cytokinetic defects in Rab11DN expressing gonads (Fig.S7B & E), confirming important roles of Rab11 mediated membrane trafficking during cytokinesis.

Rab7-null mutant flies generated by imprecise excision of P-element resulted in a 1025 bp deletion that removed most of the protein coding exon and the 5’ untranslated region (UTR) [53]. The mutants displayed late pupal lethality and western blotting with Rab7 antibody confirmed the absence of Rab7 protein in the mutant pupae (Fig.S7C). To investigate meiotic cytokinetic defects in Rab7 mutants we performed testis squash preparations followed by phase contrast microscopy. Interestingly, we found significant increase (15%) in meiotic cytokinesis defects in Rab7 mutants (Fig.S7D & E) compared to wild type controls where cytokinetic defects are virtually zero [54, 55] (Fig.S7A), suggesting a significant role for Rab7 in male meiotic cytokinesis. Previous studies have shown that Rab7 protein is required downstream of late endosomes or multivesicular bodies (MVBs) for transfer of cargo to lysosomes [56], therefore formation and migration of late endosomes to the cleavage furrow may not be affected in Rab7 mutants. Although less efficient, we observed that PlyA2-mCherry continued to be endocytosed and targeted to cleavage furrow in Rab7 mutants (Movie S13). The partial cytokinetic defects in Rab7 mutants indicates existence of alternative mechanisms to compensate for loss of Rab7 during cytokinesis. Future experiments with other Rab protein mutants in combination with Rab7 mutants might help address this possibility. The lethal nature of Rab7 mutants during the pupal stage and the time required for inducing Rab11DN during pupal stage posed technical difficulties in creating double mutants and measuring cytokinetic defects in these backgrounds.

### Multivesicular bodies deliver CPE laden vesicles to the cleavage furrow

We performed correlative light and focused ion beam scanning electron microscopy (CLEM/FIB-SEM) to gain structural insight into the endosomes that are enriched at the cleavage furrow/midbody during cytokinesis. Spermatocytes expressing EYFP-Rab7 were labeled for endocytosed CPE with PlyA2-mCherry and allowed to progress to meiosis 1. Cysts showing PlyA2-mCherry and EYFP-Rab7 co-localization at the cytokinetic furrow were fixed, imaged by light microscopy, then stained, resin embedded and prepared for FIB-SEM. Figure 7A-C shows CLEM-FIB-SEM of a cyst undergoing cytokinesis showing colocalization of Rab7 with endocytosed CPE at the furrow. 3D correlation of the LM and FIB-SEM image volumes allowed us to localize the Rab7/PlyA2 double positive signals to endosomes that appeared as spherical shaped structures that ranged from 300 to 800 nm in diameter. The lumen of these endosomes was filled with intraluminal material which at FIB-SEM resolutions could be discerned, but not definitely resolved, as vesicles packed into the lumenal volume. Although we were able to capture endosomes in the cytokinetic furrow it was technically difficult to visualize and arrest them in physical association with the ingressing furrow membranes using these flies (Movie S14). Therefore, we decided to examine cytokinesis in cysts where the contractile ring in proximity to the ingressing membrane and the endosomes could be visualized during live imaging.

**Fig. 7.**
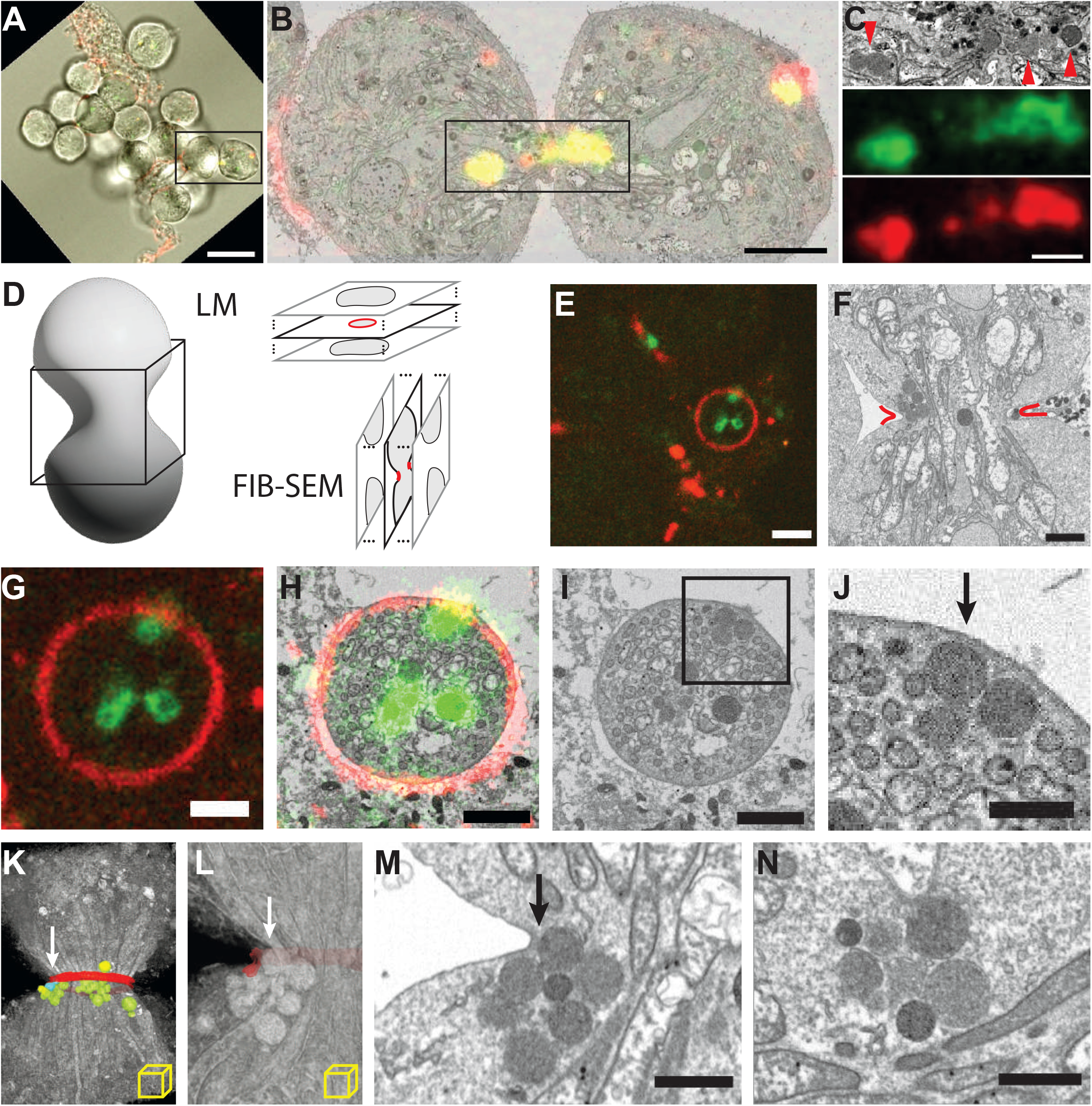
CPE positive endosomes dock at the ingressing cytokinetic furrow during meiosis. A. Combined transmitted and fluorescence image of dissected Drosophila cyst; dividing cells chosen for FIB-SEM imaging are boxed. Green, eYFP-Rab7; Red, PlyA2-mCherry. Scale bar 20 µm. B. 2D section from FIB-SEM reconstruction correlated with “z slice” from confocal. Colors as in A, saturated for ease of visualization. Scale bar 5 µm. C. Area boxed in B; FIB-SEM image (top), Rab7 (middle), PlyA2 (bottom). Arrowheads indicate electron-dense endosomes. Scale bar 2 µm. D. Volumetric imaging of dividing spermatocytes, showing orthogonal imaging planes, and expected cross sections for light microscopy (LM) and FIB-SEM. E. Fluorescence “z slice” capturing a cleavage furrow in-plane. Scale bar 4 µm F. FIB-SEM image capturing same furrow as in E cross-section, approximated by red lines. Scale bar 2 µm. G, H, I. Correlated fluorescence image, overlay, and rotated FIB-SEM section, respectively, of furrow in-plane. J. Area boxed in I showing endosomes docked at the furrow (arrow). Scale bar 1 µm. J. K. Volume rendering of FIB-SEM 3D reconstruction, with furrow segmented and false colored red, undocked endosomes green, docked endosomes blue. L. Close-up of K, with furrow segmented translucent and endosomes unsegmented M. Matching FIB-SEM section in the imaging plane. Arrows in K, L, M show same endosomes docked at furrow. Scale bar 2 µm. N. Independent CLEM/FIB-SEM experiment revealing at higher pixel sampling another example of docked vesicle. Scale bar 1 µm.

We examined spermatocytes expressing tubulin-Gal4>mRFP-Anillin that labels the contractile ring and endogenously expressing EYFP-Rab7 that marks late endosomes to capture membrane proximal endosomes (Movie S15). Anillin (also known as scraps) encodes for a conserved pleckstrin homology domain containing protein that was shown to bind actin, microtubules and nonmuscle myosin II and required for stabilization of contractile ring [39]. Dividing cysts were imaged, fixed and cysts with Rab7-Anilin colocalization at the cytokinetic furrow were prepared for FIB-SEM imaging (Fig. 7D and E). Using CLEM/FIB-SEM we were able to capture multiple Rab7 positive endosomes docked at the ingressing furrow (Fig. 7E-M). Movie S16 shows the 3D reconstruction of the vesicles captured at the furrow seen in Fig. 7K. We observed multiple instances of this docking of endosomes in dividing spermatocytes (Fig. 7N). Movie S17 is a sub-volume reconstruction of the dividing cells seen in Figure 7N. While we observed docking of the endosomes to the ingressing membrane in the samples we studied, we were unable to visualize the fusion of vesicles to these ingressing membrane or to the proteinaceous structure that was anchoring the endosomes (Fig. 7 and Movie S17). To visualize the endosomes at the furrow at higher resolution, we carried out CLEM-SEM-TEM of the dividing meiotic spermatocytes and examined vesicles that appeared to be in close proximity to the cleavage furrow (Fig.8). *Drosophila* spermatocyte cysts expressing tubulin-Gal4>mRFP-Anillin and EYFP-Rab7 undergoing meiotic cytokinesis were identified, fixed and resin embedded. Serial sectioning of the correlated cysts followed by TEM imaging revealed the structures as multivesicular body like organelles, filled with intraluminal vesicles varying in size between 20-50 nm in diameter (Fig. 8C and D1), consistent with previously described multivesicular endosomes (MVEs) or Multivesicular bodies (MVBs) [57]. Again, rather than seeing evidence of fusion, we found that outer membrane of several of these endosomes was discontinuous (Fig.8C). Interestingly, the MVBs that showed significant loss of outer membrane integrity appeared to have released intraluminal vesicles in the vicinity of the ingressing membranes (Fig. 8D, top left), possibly providing a proximal source of lipid laden vesicles for membrane biogenesis (Fig. 8E).

**Fig. 8.**
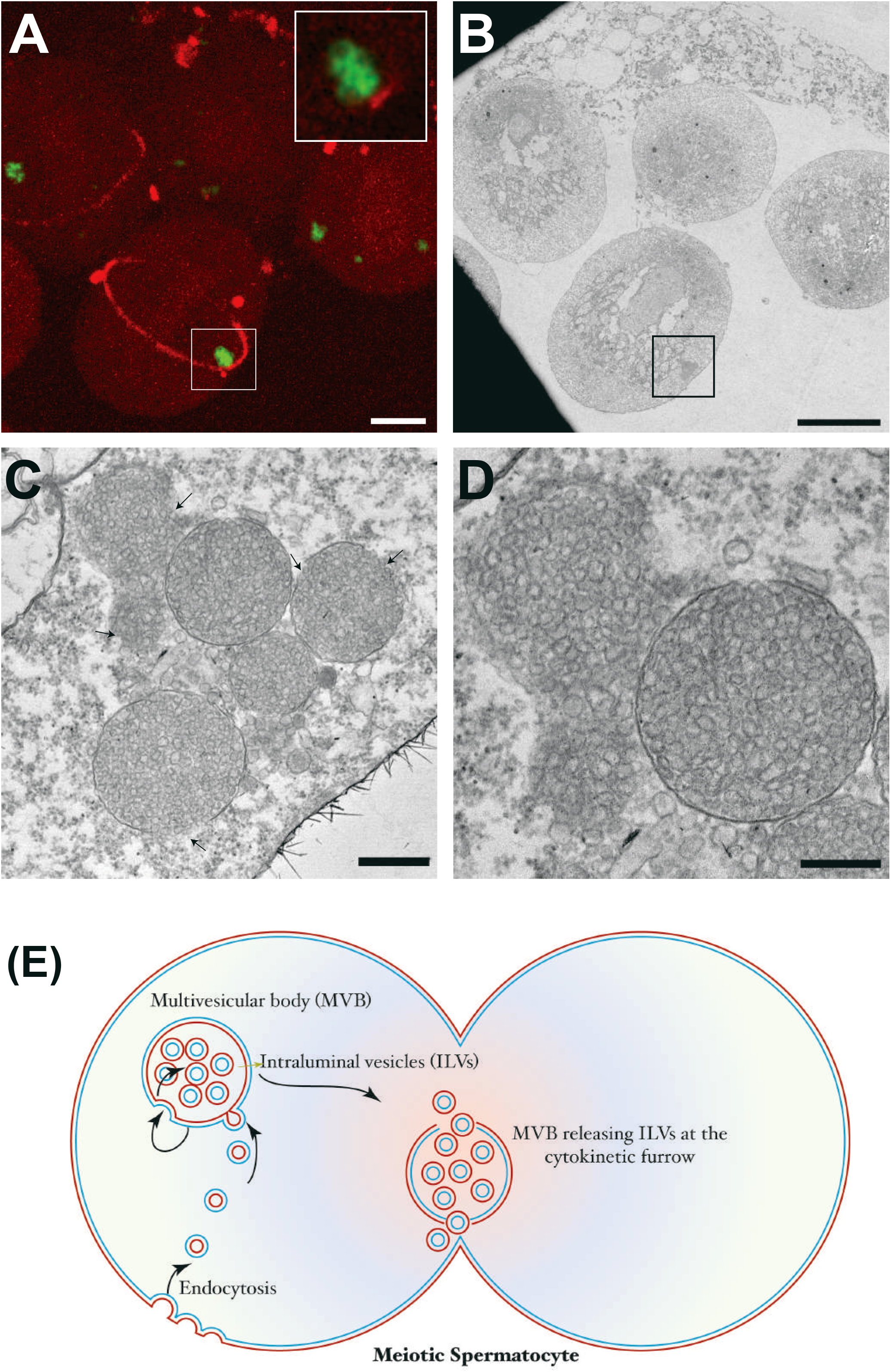
Multivesicular bodies release intraluminal vesicles at the cytokinetic furrow. A. Fluorescence maximum projection image capturing dividing cells from a dissected drosophila cyst. Green, eYFP-Rab7; Red, mRFP-Anillin. Inset, vesicles adjacent to a cleavage furrow. Scale bar 10 µm. B. Corresponding TEM section. Endosomes adjacent to the cleavage furrow are boxed. Scale bar 10 µm. C. Higher magnification TEM image of area boxed in B. Scale bar 500 nm. Arrows indicate discontinuity in outer membranes of MVBs. D. Subsequent TEM section of same area, showing make-up of vesicles at high magnification. Scale bar 300 nm. E. A model depicting endocytosis of CPE from plasma membrane, the maturation of these endosomes into multivesicular bodies, targeting and release of intraluminal vesicles at the cleavage furrow.

## Discussion

Plasma membrane expansion and ingression of the cytokinetic furrow still remain one of the least understood horizons in the field of cell division [58]. We are only beginning to appreciate the involvement of lipids in structural integrity, membrane expansion, and transmission of signals during cytokinesis. PIP2 is the most studied lipid in cytokinesis and has been shown to localize to the cleavage furrow during ingression. In addition to PIP2, cholesterol, very long chain fatty acids and sphingolipids have also been associated with cytokinesis [59] [10, 12]. In mitotic HeLa cells, PIP2 accumulation at the inner leaflet and subsequent recruitment of RhoGTPase (RhoA) was shown to be coordinated with sphingomyelin rich domains on the outer leaflet [10]. However, it is unknown if a similar mechanism exists in germ cells which have different characteristics and different lipid composition. While mitotic cells were shown to accumulate classical sphingolipids with long to very long acyl chains at intercellular bridges [12], male germ cells accumulated complex sphingolipids with very long/ultra long chain polyunsaturared fatty acids (VLC-PUFAs) [13]. These differences in sphingolipid composition with distinct biophysical properties between mitotic cells and male meiotic cells point towards existence of distinct underlying mechanisms.

In this study, we show the sphingolipid composition of the major species in *Drosophila* testis for the first time and demonstrate significant enrichment of unique CPE species containing SPH_MUFA and SPD_SFA. Unlike mammalian testis, the *Drosophila* testis lacks significant amounts of VLC-PUFA containing sphingolipids. However, due to conserved unsaturation in acyl chains, it is likely that SPH_MUFA and SPD_SFA containing CPE could substitute for VLC-PUFA functions in *Drosophila* male germ cells. Glycosphingolipids with VLC/ULC PUFAs have been shown to be required for mammalian male meiotic cytokinesis but their actual role has not been delineated [13, 60, 61]. It is interesting to note that CPE mimics certain physical properties of galactosylceramide (GalCer) a major sphingolipid in myelin sheath [62]. Previous studies have shown that CPE has an essential role in axonal ensheathment and cortex glial plasma membrane expansion in *Drosophila* [15, 63]. It was hypothesized that the axonal ensheathment in mammals and *Drosophila* is based on similar physical process with different lipids [62]. We have previously shown that sphingomyelin, a structural analog of CPE, could rescue cortex glial membrane defects in *cpes* mutants suggesting the head group is not as important as the tail in glial membranes. However, here we show that sphingomyelin could not rescue spermatogenesis defects in *cpes* mutants alluding to the importance of the ethanolamine head group in spermatocytes. Taken together, we could suggest that *Drosophila* CPE would have evolved to have both the properties of mammalian glycosphingolipids and sphingomyelin to promote spermatogenesis and glial ensheathment of neurons.

Phosphoinositides play critical role in cytokinesis. In particular, PIP2 was shown to be enriched at the furrows and directly binds to several proteins such as anillin, septins and MgcRacGAP to mediate furrow stability [64, 65]. Unlike mitotic cells, PIP2 in male meiotic cells does not show selective accumulation at cleavage furrows. Instead, it is uniformly distributed throughout the plasma membrane including at furrows ([45] and Fig.4C and Movie S5, S6 and S7). Interestingly, PIP2 hydrolysis and calcium release were shown to be required for cytokinesis in *Drosophila* spermatocytes [45]. These observations suggests that in meiotic cells, PIP2, has a broader role at the plasma membrane and at furrows to promote cytokinesis. Intriguingly, we show that the PIP3 binding PH domain of Steppke (tGPH) localizes to cleavage furrows indicating a potential role for PIP3 during male meiotic cytokinesis. However, the accumulation of PIP3 to the cleavage furrows occurs only in the intact cysts i.e, 16 interconnected spermatocytes wrapped up by two somatic cyst cells suggesting the need for certain mechanical force. The role of PIP3 accumulation in the male meiotic cleavage furrows is an open question and warrants future investigation. Notably, recently it was shown that cytohesin Steppke reduces tissue tension by inhibiting the actomyosin activity at adherens junctions in embryos [46]. Although broad accumulation of PIP2 to the plasma membranes and PIP3 accumulation at the cleavage furrows was not altered in *cpes* mutants, their specific localization to membrane microdomains and their inability to signal robustly cannot be ruled out. Sphingadienes with *Δ*^4,6^ conjugated double bonds were previously identified in *Drosophila*, *Manduca sexta*, and *Bombyx mori* [66–68], although their enrichment in the testis was not known until now. However, normal sphingolipid synthesis and their degradation are shown to be required for spermatogenesis [69, 70]. The sphingadiene sphingolipids were shown to have inhibitory effects on AKT-dependent signaling [71] and downregulate *Wnt* signaling via a PP2A/Akt/GSK3*β* pathway [72]. Thus, accumulation of sphingadiene with saturated fatty acid containing sphingolipids in the testis could have a role in regulation of PIP2/PIP3 mediated signaling *in vivo*. Apart from CPE_SPD_SFA the *Drosophila* testis is also enriched in CPE_SPH_MUFA suggesting they could mediate distinct functions. Lipids with unsaturated fatty acids mediate a number of biological functions [73]. It is widely known that phospholipids with unsaturated fatty acids promote membrane fluidity and elasticity [74, 75]. Sphingolipids with unsaturated fatty acids could thus provide unique biophysical properties necessary for the membrane curvature requirements during meiotic cytokinesis. *des-1* codes for dihdryoceramide desaturase enzyme in the *de novo* biosynthetic pathway in *Drosophila*. *des-1* mutants (also called *infertile crescent*, *ifc*) were described before the gene product was identified as a homolog of dihydroceramide desaturase. The primary spermatocytes of the *des-1* mutants undergo degeneration without initiating chromosome condensation at the beginning of meiosis [76]. A P-element insertion allele showed that DES-1 colocalized with mitochondria and was intimately associated with the central spindle of dividing meiotic spermatocytes. Its deficiency leads to failure of central spindle assembly in these dividing cells leading to several phenotypes including cytokinetic defects. It was proposed that DES-1 could be part of an anchoring mechanism that linked membrane bound cellular compartments to components of the cytoskeleton [69]. Considering that DES-1 is a dihydroceramide desaturase homolog, it would be worth revisiting the mutant phenotype to evaluate if it too contributes to membrane assembly at the cytokinetic furrow due to lack of CPEs.

In correlative live imaging and electron microscopy studies, we find that multivesicular bodies (MVBs) target CPE laden vesicles to the cleavage furrow. MVBs are by definition spherical organelles that are typically 250-1000 nm in diameter with a single outer membrane that enclose a variable number of smaller spherical vesicles within. Originally they were believed to be involved in neurosecretion because of their close association with the Golgi stacks [77]. Further work established a primary role in endocytosis and subsequent degradation of proteins [78–80]. Other work have highlighted the important role of MVBs in recycling function [81]. The last decade or so has seen the emergence of a role for MVBs in autophagy, and secretion via the exosomes [82, 83]. Our work has surprisingly unraveled a possible new function for MVBs in transporting the endocytosed vesicles rich in lipids to the vicinity of expanding membrane. Interestingly, CPE is not required for localization of MVBs to the cleavage furrow (Movie S18), instead, our data suggests that CPE loaded MVBs are required in the vicinity of expanding membranes to mediate cytokinesis. Future studies should throw light on the role of this pathway in membrane and protein accumulation in newly synthesized membranes as in the case of meiotic cytokinesis in the testis. The human genome encodes more than 60 Rab proteins and several of them including Rab1, Rab8, Rab10, Rab14, Rab21, Rab24 and Rab35 were shown to localize to intercellular bridges. However, most of the studies primarily focused on the recycling role of Rab11 and Rab35 endosomes and their effector proteins in cytokinesis [65]. Currently it is unknown if these endosomes are also involved in delivery of specific plasma membrane derived lipids to the cleavage furrows. Our study shows that there is an increased localization of CPE to the endosomal compartments in spermatocytes. The CPE containing MVBs are selectively docked on the ingressing membranes at the cytokinetic furrow and release intraluminal vesicles, suggesting delivery of CPE rich membranes to the growing furrows. Future investigations focusing on these CPE enriched endosomes and their dyamics during male meiotic cytokinesis, spermatid polarity and individualization will provide novel insights into membrane expansion and or their role as a signaling center to mediate spermatogenesis.

## Materials and Methods

### Fly stocks

tubulin-Gal4 (BDSC#5138), UAS-PLC*δ*-PH-EGFP (BDSC#39693), tGPH (BDSC#8163), Sqh-GFP.RLC (BDSC#57145), Sqh-mCherry.M (BDSC#59024), p(Ubi-p63E-Feo-mCherry)3 (BDSC#59277), *bam*-Gal4 [37], *nos*-Gal4 (BDSC#4937), *chif*-Gal4 (BDSC#13134), C587-Gal4 (BDSC#67747), EYFP-Rab4 (BDSC#62542), EYFP-Rab6 (BDSC#62544), EYFP-Rab7 (BDSC#62545), EYFP-Rab11(BDSC#62549), UAS-EGFP (BDSC#5430), tub-Gal80ts (BDSC#7108), UASP-Rab11.S25N (BDSC#23261), UAS-mRFP-Anillin (BDSC#52220), rab7 mutant (Krisztina Hegedus et al., 2016), genomic CPES_V5tag [15], *cpes* mutants [15], UAS-hSMS1 [15], dcert1[31], UAS CDase [32], UAS-CPES active site mutants, UAS-Aedes CPES, UAS-Bombyx CPES (this study).

### Fly husbandry

All *Drosophila* stocks were raised on standard fly food and maintained 25°C unless otherwise mentioned. Due to increased lethality *cpes* mutants were raised at 18-21°C. Certain genetic backgrounds increased pupal death in *cpes* mutants such as overexpression of UAS CDase, UAS-PLC*δ*-PH-EGFP, *cpes*;*dcert^1^* double mutants. In such cases we have separated 3^rd^ instar larvae of *cpes* mutants and allowed them to develop into pupae. The testes were dissected before the death of pupae and analyzed.

### Cloning of UAS-*Aedes egypty* CPES and UAS-*Bombyx Mori* CPES

The coding sequence for *Aedes aegypti* CPES <u>(XM_021851518.1</u>), and *Bombyx mori* CPES (<u>XM_004923146.3</u>) were codon optimized and synthesized by GenScript® and subcloned into pUAST vector. The clones were sent to BestGene Inc for embryo injection service. The transgenic flies were balanced and crossed with Gal4 drivers to conduct rescue experiments.

### Cloning of CPES active site mutant

Two conserved aspartates in CPES active site were mutated to alanine using PCR based site directed mutagenesis method as described by [84]. PCR was performed using pBluescript SK+ CPES_V5 clone as template with appropriate sense and antisense primers (Table S1) and Phusion polymerase (NEB). The PCR product was digested with DpnI to remove templates followed by transformation into DH5*α* competent cells. Plasmids were isolated from colonies grown in LB media and screened by restriction digestion with appropriate restriction enzymes (Table S1) and sequencing. The pBluescript SK+ CPES DD-AA clone was used as template to PCR amplify the insert using appropriate primers (Table S1), restriction digested and cloned into pUAST vector. The clones were confirmed with restriction digestion followed by DNA sequencing.

### Cloning, expression, and purification of PlyA2mCherry

Mushroom-derived aegerolysin PlyA2 was cloned, expressed, and purified as described by [85]. Briefly, the coding sequence of PlyA2 (accession number AB777517) fused to mCherry at the C-terminus was synthesized by GenScript®. The linker amino acid sequence between PlyA2 and mCherry was VDGTAGPGSIAT. This clone was used as template to PCR amplify using appropriate forward and reverse primers (Table S1) and cloned into p24a vector at appropriate restriction sites (Table S1). The clones were transformed into Escherichia coli strain BL-21 (DE3) and single colony was inoculated in to 5ml of LB broth containing 50μg/ml kanamycin and grown overnight in orbital shaker (200rpm) at 37°C. 2ml of overnight culture was inoculated into 250ml of fresh LB broth containing kanamycin and grown at 37°C until the optical density reached to 0.6. Subsequently, the culture was incubated in cold (4°C) for 2 hours, followed by induction with 0.3mM isopropyl-B-D-thiogalactopyranoside and grown at 25°C in orbital shaker (200rpm) overnight. The cells were harvested by centrifugation at 5,000g for 10min at 4°C and the pellet was resuspended in binding buffer (20mM sodium phosphate buffer (pH 7.4), 500mM NaCl, 20mM imidazole) containing EDTA free protease inhibitor cocktail (Sigma, P8340). The cell suspension (50 ml) was sonicated at 4°C (Cole-Parmer ultrasonic processor model CP130,) with microprobe at an amplitude of 30 and on and off cycles of 10sec each for about 30 min to 1 hour. The lysate was centrifuged at 10,000g for 10min to remove insoluble proteins and debris. To the supernatant, 1ml of 50% Ni-NTA agarose (Thermo Scientific) was added and incubated on end-to-end rotor for 3 hours and the beads were collected on a column and washed with 30ml of binding buffer and eluted with binding buffer containing 200mM imidazole. The fractions were dialysed in 20mM sodium phosphate buffer 3 times (1L each) and the final protein was concentrated with Amicon centriprep filter devices for volumes up to 15ml. The protein concentration was determined using Bradford assay. The proteins were resuspended in 20% glycerol, aliquoted (100μl) and stored at −80°C for future use.

### Testis squash preparation, phase contrast microscopy, and immunofluorescence microsocpy

*Drosophila* testis squash preparations for phase-contrast microscopy and immunostaining were performed as described by [17]. Briefly, young male (0-2 days old) testis were dissected (5 pairs) in phosphate-buffered saline (PBS, 130 mM NaCl, 7mM Na2HPO4, 3mMH2PO4). The testes were torn open on a slide at appropriate location with a pair of Black Anodized Steel needles (tip diameter 0.0175mm) and a cover glass was gently placed without air bubbles. The slides were directly observed under phase-contrast microscopy or snap frozen in liquid nitrogen for immunostaining. After freezing the cover glass was removed from the slide with razor blade and immersed in cold 99.5% ethanol and incubated in −20°C for 10 min. The slides were then fixed with 4% paraformaldehyde (PFA) in PBS containing 0.1% Triton-X-100 (PBST) for 15 min. Subsequently, the slides were washed with PBS and the area of squashed tissue was circled with a hydrophobic barrier pen, permeabilized with PBST (30 min) and blocked with 5% normal goat serum in PBST (1 hour at RT). Squashed tissues were incubated with primary (1μg in 100μl) and secondary antibody (1:100 dilution) (8-12 hours each at 4°C in a moist chamber). Slides were washed with PBST (3x 10 min each) after each antibody treatment, finally incubated with DAPI solution (1:2000 dilution from 5mg/ml stock, Molecular Probes) and mounted with Vectashield H1000 mounting media.

### Quantification of cytokinetic defects

5 pairs of testes were dissected for each experiment, live testis squashes were prepared and round spermatids were observed using 40X Ph2 Plan-Neofluar objectives on Axioplan 2 microscope as described above. All round spermatid cyst in the sample were imaged and their cell number was counted manually (250-500) for each genotype. Percentage of round spermatids with 2, 3 or 4 nuclei were calculated by dividing the with total spermatid number including spermatids with 1, 2, 3, or 4 nuclei. More than 5 independent experiments were carried out for each genotype.

### Male fertility assay

Male fertility assay was performed by setting up individual crosses with 1 male and 3 wild type females. Each replicate involved 30 males and three independent replicates (total of 90 males) were performed for each experiment.

### Immunostaining and confocal imaging

*Drosophila* tissues were dissected and immunostained as described previously [15]. Briefly, dissected testes were fixed with 4% PFA in 0.3% PBST for 1 hours at RT, washed (3x 10 min each), permeabilized in 0.3% PBST (1 hour), blocked with 5% NGS (1hour) and incubated with primary antibody (1:100 dilution or 1μg in 100μl of 5% NGS in PBST) overnight (8-12 hours). Testes were washed with 0.3% PBST (3×10min each) and incubated with secondary antibody (1:100 dilution in 5%NGS in PBST). Testes were washed with PBST (3x 10 min each), stained with DAPI (1:2000 dilution of 5mg/ml solution in PBST) for 15 min, and mounted with Vectashield H1000 mounting medium. The slides were imaged on ZEISS confocal laser scanning microscope LSM 780 or LSM880.

### Live cell imaging of meiotic cysts

About 15-20 pairs of young (0-2days old) *Drosophila* testes were dissected in 600μl of insect cell culture media (M3 media supplemented with heat inactivated FBS, Pen/Strep) in a glass well plate. Testes were transferred to a fresh well with 400μl of media and torn open with a pair of Black Anodized Steel needles (tip diameter 0.01mm) to release the cysts into media. This was repeated until all the testes were torn open. They were gently agitated so that all the cysts were released into the media followed by removal of empty testis or debris from the media. The cysts were gently washed with fresh 400μl media (2x), then transferred *via* pipetting into the middle of 35mm cover glass bottom dish (Poly D-Lysine coated) and fresh media (max. 200μl) was gently added to the cleft formed between cover glass and the dish. The cysts tend to move while handling but remain settled on the cover glass and ideally remain in position while imaging. The intact cysts and cysts that have lost their cyst cells could be distinguished in brightfield view. The cysts that are beginning to undergo meiosis could be identified in brightfield view by looking at cell morphology and differences in nucleus and spindle appearance. Cells beginning to undergo meiosis could also be identified by Feo-mCherry which is localized to nucleus in interphase but translocates to cytosol during prophase to anaphase. Prophase to anaphase cysts were identified using 25x water objective and subsequently imaged using 63x water objective on a Leica Andor spinning disk confocal microscope. The time lapse imaging was done at 2 min intervals with the z stacks (about 80 slices, 0.5μm per slice) at each time point for 60 to 90 min for completion of meiotic I cytokinesis. Laser power and exposure times were preferably kept low to prevent bleaching and aberrant effects on cytokinesis due to generation of free radicals.

### Sample preparation for correlative light and electron microscopy (CLEM)

About 10-15 pairs of testes were dissected from newly eclosed male flies in M3 or S2 insect cell culture media. Testes were transferred to fresh M3 media (400 µl) in a Glass Spot plate (PYREX 722085) and torn open with a pair of black anodized steel needles and gently agitated to release the cysts into media. Subsequently, testis debris were removed from the well. The cysts were gently washed with fresh 400 µl media (2x) and incubated with PlyA2-mCherry protein (10μg /ml) in 400μl of M3 media for 1 hour, on a rocker. The cysts were washed with fresh media without PlyA2-mCherry (1x with 400μl media). About 5-10 cysts were transferred to cover glass bottom 96 well plate (Cellvis, Cat # P96-0-N) containing 100μl of M3 media. The cysts in each well were observed under 25X water objective on an Andor spinning disk microscope using bright field and or fluorescence. Cells undergoing division were identified by cell morphology changes such as cell rounding, nuclear membrane loss, microtubule appearance, cell elongation and ingressing cleavage furrows. When spermatocytes at the appropriate stage of interest were identified, 100 µl of 8% PFA in M3 media was added to the well (4% PFA final concentration) and gently mixed with pipetting. The 96 well plate cover was removed after adding PFA to avoid its effect on other wells. Location of the well could be identified with the help of a torch light and or illuminating the well with bright field or fluorescent light. The process was repeated until at least 5-10 cysts were identified, and fixed. The sample in fixative could be stored up to 2 days at 4°C in the 96 well plate. The cysts in the 96 well plate were gently agitated with pipetting and transferred to a glass spot plate (the pipet tip was cut and conditioned with the media to avoid sticking of cysts to tip walls). At this stage cysts were pooled from different 96 plate wells into a single glass well plate. The cysts were centered by gently flushing with pipetting. They were then washed with 1XPBS (3X 400 µl). Following washing, the cysts were pipetted into the center of a gridded covered glass bottom plate (MatTek, Part No. P35G-1.5-14-C-Grid). After the cysts settled on the coverslip, most of the PBS was removed by gentle pipetting. Subsequently 4% low melting agarose (Invitrogen, REF 16520-100) (in 1xPBS) that was maintained at 80 °C is cooled for 1-2 min and gently poured from the top, care must be taken to minimize movement of cysts. After the agarose had solidified, the cysts that were settled and embedded in the agarose were imaged on the Andor spinning disk confocal microscope using 25X and 63X water objectives. At this time, the cells were imaged at high magnification by transmitted light to record the location of regions of interest (ROI) with respect to the alphanumeric grid of the glass-bottom dish. Images were also taken at lower magnification to record the locations of other cells in the dish, which would later serve as landmarks in the correlative light-electron microscopy (CLEM) pipeline; images of the ROI at various points in this pipeline are shown in Fig.S8.

The disc of solidified agarose containing the cysts was carefully removed from the glass-bottom dish. Not only are the cells embedded in situ typically within 10 microns of the bottom agarose surface, but the alphanumeric gridded pattern is also transferred to the agarose pad in relief, and both are visible in a dissection microscope. The agarose pad was rinsed with 0.1 M sodium cacodylate buffer (11652, Electron Microscopy Sciences) several times. It was then incubated in a solution containing 2% osmium tetroxide (19150, Electron Microscopy Sciences) and 1.5% potassium ferricyanide (20150, Electron Microscopy Sciences) in 0.1M sodium cacodylate buffer for 1 hour at room temperature. Afterwards, it was thoroughly washed with water for a minimum of 30 minutes. The sample was then incubated in 1% aqueous uranyl acetate (22400, Electron Microscopy Sciences) for 30 minutes followed by extensive water washes. This was followed by standard EM processing steps – the sample was dehydrated using a series of increasing ethanol concentrations ending in 100% ethanol, and then washed with 100% propylene oxide. The sample was infiltrated with Epon (Embed 812) resin using increasing concentrations of hard resin formulation in propylene oxide, and finally embedment in a polypropylene dish, with curing in an oven at 60C for 2 days. Once the resin was fully cured, it was separated from the dish and examined using a dissecting microscope. The heavy metal-stained cysts were dark and easily identified; the overall pattern of cysts was unchanged from previous steps (Fig.S8). ROIs for CLEM were identified and ∼1 mm^2^ areas around each cyst were cut out from the larger disc of resin using a saw and razor blades. These were then re-embedded onto blank blocks for ultramicrotomy. The blocks were sectioned using an ultramicrotome (Powertome, RMC) until cysts were exposed at the resin surface as revealed by Toluidine Blue stain, taking care that the actual cleavage furrow itself was just below the surface and not sectioned. This positions the feature within ∼10 microns of the top surface and therefore amenable to FIB-SEM imaging.

### FIB-SEM, STEM and TEM imaging

FIB-SEM imaging was done largely as previously described [86]. Briefly, the specimen was cut to ∼ 2mm height and mounted on a SEM stub using silver. The specimen was introduced into a Zeiss CrossBeam 550 (Zeiss Inc.), ROI located by SEM imaging, and protected by a patterned platinum and carbon pad. A trench was FIB milled until the profiles of both dividing cells were revealed but stopped before the cleavage furrow was milled away. Images were acquired at 3 nm or 5 nm pixel sampling in XY, and step size of 9 or 15 nm in Z, respectively, in an automated FIB-mill-SEM-image cycle with SEM operated at 1.5 kV, 1.2 nA; FIB milling at 30 kV, 1.5 nA, and back-scatter detector grid voltage at 900 V. The image stack was registered, contrast inverted and binned by 3 in the imaging plane to produce an isotropic 9 nm^3^ or 15 nm^3^ image volume reconstruction. For TEM imaging, the cysts were exposed using the same approach, but here, 60 nm serial sections were cut and collected onto TEM grids. STEM imaging was executed at a Zeiss Gemini II SEM 450 at 30 kV landing energy, and the TEM imaging was done on a Hitachi 1050 operated at 80 kV. In both cases, the grids were not post-stained, however a ∼4nm carbon coat was applied using a sputter coater (Leica Inc.) before imaging.

### Image correlation and visualization

There were two stages of correlation in these experiments. The correlation for relocation, i.e., imaging an ROI by LM and then locating the same ROI for EM, was performed by careful sample preparation and appropriate imaging at various stages (Supp Fig S1). Correlation for registration, i.e., aligning two image volumes to identify cellular features of interest, was done after LM and EM images were acquired. 3D correlation was done using eC-CLEM [87]. The confocal stack and FIB-SEM image stacks were imported, and multiple fiducials were placed at distributed locations on the plasma membrane of both dividing cells in the PlyA2-mCherry + eYFP-Rab7 expressing cysts. The plasma membrane was visualized easily in the FIB-SEM images, and more approximately in the lower resolution LM data, using the PlyA2-mCherry signal which localizes strongly at the cell membrane. The green channel (Rab7) was not used for registration. The skew transformed LM fluorescence was overlaid on the original FIB-SEM data and examined with the green channel added back, upon which the identity of the PlyA2 + Rab7 double positive features in the furrow were revealed (Supplementary Movie S14). In a second experiment using mRFP-Anillin:eYFP-Rab7 cysts (Fig. 7E), the correlation in ecCLEM was performed using fiducial points on the furrow, as located by the “neck” in the FIB-SEM image volume (Fig. 7F), and the mRFP-Anillin staining in the LM images (Fig. 7G). As before, the Rab7 signal was not used for correlation. In both cases, the registration was limited by the relatively lower resolution of the LM image stack but was sufficient to consistently correlate large vesicles in the FIB-SEM data with Rab7-positive signals in the LM data. For TEM imaging, sequential sections were imaged and inspected manually until the features captured in the LM started to appear in the EM sections. The images were overlaid but not computationally correlated, as the identity of the features was readily apparent in the 2D EM image. The FIB-SEM image volumes were visualized, and volume rendered using Arivis (Arivis Inc), with features of interest such as furrow, docked and undocked vesicles segmented either in 3DSlicer (www.slicer.org) or Arivis.

### Sphingolipid estimation in fly tissues using supercritical fluid chromatography coupled to mass spectrometry (SFC/MS/MS)

Sphingolipid estimation using supercritical fluid chromatography coupled to mass spectrometry (SFC/MS/MS) was performed as described previously [15]. Lipids were extracted from heads, ovary and testis samples as described [88, 89]. About 400 fly heads, about 250 pairs of ovaries and about 250 pairs of testes were used for each biological replicate. Three independent replicates were taken for each sample. For Figs.1 and 2 only testes samples were used whereas for Fig.4 whole male organ (testis, accessory glands, collecting ducts and ejaculation bulb) were used for lipid extraction. Internal standards (ISTD) added to the tissues before extraction included 500 pmol (20 μl) of Cer/Sph Mixture I (Avanti Polar Lipids LM-6002), and 1nmol of C12 Sphingosyl PE (C17:1/C12:0) (Avanti Polar Lipids). Lipids from all samples were normalized to carbon content (100μg/ml) and 100μg/ml sample was diluted 10 times before injection for SFC. We have used large-fragment MRM method for quantitation of amount per 100μg of carbon.

### Total RNA sequencing

Three independent biological replicates were performed for each sample and about 200 pairs of *w^1118^* (WT 1-3), *cpes* mutant (KO 1-3) and *bam*-Gal4>UAS-CPES rescued (RES 1-3) fly testis were dissected for each replicate. The testes were dissected in S2 cell culture media, washed with PBS and RNA was extracted. Total RNA was extracted using Trizol method followed by DNAase I treatment and purification using RNA Clean and Concentrator^TM^-5 kit (Zymoresearch). A total of 2μg of RNA was submitted to Novogene for total RNA seq. The RNA seq data was further analyzed for differential gene expression and pathway analysis using Gene Set Enrichment Analysis (GSEA). The gene sets specific for *Drosophila* were downloaded from http://www.bioinformatics.org/go2msig/ and http://ge-lab.org/gskb/. The gene sets specific for cell types in *Drosophila* testis are described in Evan Witt et al., 2019 [33].

## Acknowledgments

This study was funded by the intramural division of the National Cancer Institute, National Institutes of Health (Division of Health and Human Services). The content of this publication does not necessarily reflect the views or policies of the Department of Health and Human Services, nor does mention of trade names, commercial products, or organizations imply endorsement by the U.S. Government. The study was partly funded with Frederal funds from the National Cancer Institute, National Institutes of Health, under Contract No. 75N91019D00024. The work was partially supported by the Grant-in-Aid for Scientific Research on Innovative Areas (17H06304) [T.B.] and a Grant-in-Aid for Scientific Research (B) (18H01800) [T.B.] from Japan Society for the Promotion of Science (JSPS). We thank Mr. Aayush Bhtawadekar for help with movies and rendering for the FIB-SEM images.

## Conflict of Interest Statement

The authors declare no conflicts of interests.

## Supplemental movie legends

**Movie S1.** Live cell imaging of neuroblast asymmetric divisions in 3^rd^ instar larval brain. GFP tagged PH domain of PIP3 binding protein Step was expressed under the control of alpha tubulin84B promoter. Arrows indicate enrichment of GFP-Step on cleavage furrow membranes during cytokinesis.

**Movie S2.** Live cell imaging of an intact control *Drosophila* spermatocyte cyst undergoing meiosis I cytokinesis. Spermatocytes are expressing myosin II regulatory light chain (RLC) tagged to GFP under its own promoter and Ubi-p63E promotor drives the expression of Feo-mCherry. An intact cyst is shown. Sqh-GFP-RLC localizes to actomyosin contractile ring. Feo-mCherry localizes to central spindle.

**Movie S3.** Live cell imaging of an intact *cpes* mutant spermatocyte cyst undergoing meiosis I cytokinesis. Mutant spermatocytes are expressing myosin II regulatory light chain (RLC) tagged to GFP under its own promoter and Feo-mCherry was expressed under the control of Ubi-p63E promotor. Sqh-GFP-RLC specifically localizes to actomyosin contractile ring. Feo-mCherry localizes to central spindle.

**Movie S4.** Live cell imaging of an intact CPES rescue spermatocyte cyst undergoing meiosis I cytokinesis. The UAS-CPES is expressed using Tub-Gal4. Spermatocytes are also expressing Sqh-GFP-RLC and Feo-mCherry. Sqh-GFP-RLC specifically localizes to actomyosin contractile ring. Feo-mCherry localizes to central spindle.

**Movie S5.** Live cell imaging of an intact control *Drosophila* spermatocyte cyst undergoing meiosis I cytokinesis. The UAS PLC*δ*-PH-EGFP is expressed using Tub-Gal4 and Feo-mCherry is expressed under the control of Ubi-p63E promoter. PLC*δ*-PH-EGFP specifically binds to PIP2 at the plasma membrane. Feo-mCherry localizes to central spindle.

**Movie S6.** Live cell imaging of an intact *cpes* mutant spermatocyte cyst undergoing meiosis I cytokinesis. Tub-Gal4 drives the expression of UAS PLC*δ*-PH-EGFP and Ubi-p63E promoter drives the expression of Feo-mCherry. PLC*δ*-PH-EGFP specifically binds to PIP2 at the plasma membrane. Feo-mCherry localizes to central spindle.

**Movie S7.** Live cell imaging of an intact CPES rescue spermatocyte cyst undergoing meiosis I cytokinesis. The UAS-CPES and UAS PLC*δ*-PH-EGFP are expressed using Tub-Gal4. The Feo-mCherry is expressed under the control of Ubi-p63E promoter. PLC*δ*-PH-EGFP specifically binds to PIP2 at the plasma membrane. Feo-mCherry localizes to central spindle.

**Movie S8.** Live cell imaging of an intact control *Drosophila* spermatocyte cyst undergoing meiosis I cytokinesis. Alpha tubulin84B promoter drives the expression of GFP tagged PH domain of Step that specifically binds to PIP3 at the plasma membrane. Feo-mCherry localizes to central spindle.

**Movie S9.** Live cell imaging of an intact *cpes* mutant spermatocyte cyst undergoing meiosis I cytokinesis. The GFP-tagged PH domain of Step is expressed under the control of alpha tubulin84B promoter and Feo-mCherry is expressed under the control of Ubi-p63E promoter. The GFP-PH-Step specifically localizes to PIP3 at the plasma membrane and Feo-mCherry localizes to central spindle.

**Movie S10.** Live cell imaging of an intact CPES rescue spermatocyte cyst undergoing meiosis I cytokinesis. PH domain of Step is tagged to GFP and expressed under the control of alpha tubulin84B promoter and Feo-mCherry is expressed under the control of Ubi-p63E promoter. The UAS-CPES expression is driven by Tub-Gal4. The GFP-PH-Step specifically localizes to PIP3 at the plasma membrane and Feo-mCherry localizes to central spindle.

**Movie S11.** Live cell imaging of EYFP-Rab7 expressing spermatocytes treated with recombinant CPE binding protein tagged to mCherry (PlyA2-mCherry). The EYFP-Rab7 is expressed under its own promotor.

**Movie S12.** Live cell imaging of EYFP-Rab11 expressing spermatocytes treated with recombinant PlyA2-mCherry. EYFP-Rab11 is expressed under its own promotor.

**Movie S13.** Live cell imaging of *rab7* mutant spermatocytes treated with recombinant CPE binding protein tagged to mCherry (PlyA2-mCherry).

**Movie S14.** FIB-SEM image stack corresponding to “Figure 7B”

**Movie S15.** Live cell imaging of control *Drosophila* spermatocyte cyst undergoing meiosis I cytokinesis. The EYFP-Rab7 is expressed under its own promotor. Tubulin Gal4 drives the expression mRFP-Anillin that localizes to contractile ring.

**Movie S16.** Movie showing the FIB-SEM volume reconstruction of the dividing cells in “Figure 1K”, highlighting the points at which vesicle docking at the furrow could be captured. Cells were 3D rendered in grayscale and overlaid by manually segmented 3D models of the cleavage furrow (red), undocked vesicles (green) and docked vesicles (blue).

**Movie S17.** Movie showing the FIB-SEM sub-volume reconstruction of the dividing cells in “Figure 8N”, highlighting the points at which vesicle docking at the furrow could be captured.

**Movie S18.** Live cell imaging of cpes mutant spermatocytes expressing EYFP-Rab7 under its own promotor and mRFP-Anillin driven by tubulin-Gal4.

---

**Fig.S1. Comparison of hexosylceramide subspecies in Drosophila tissues.** (A) Hexosylceramides composed of sphingosine base linked to saturated fatty acid (HexCer_SPH-SFA) and its subspecies are shown as picomols (pmol) in 100 micrograms of carbon. Wild type heads from male flies (WTHM), wild type heads from female (WTHF), wild type ovary (WTOY) and wild type testis (WTTS). (B) Hexosylceramide subspecies composed of sphingosine base linked to Monounsaturated fatty acid (HexCer_SPH_MUFA). (C) Hexosylceramide subspecies composed of sphingosine base linked to Polyunsaturated fatty acid (HexCer_SPH_PUFA). (D) Hexosylceramide subspecies composed of Sphingadiene base linked to Saturated fatty acid (HexCer_SPD_SFA). (E) Cartoon showing the representative chemical structure of three major and one minor HexCer subspecies including HexCer_SPH_SFA, HexCer_SPH_MUFA, HexCer_SPD_SFA and HexCer_SPH_PUFA respectively. Statistical significance was calculated using mean, standard deviation (SD) and N in Prism 8. The 2way ANOVA multiple comparison was used to calculate P values where ****P ≤ 0.0001;***P ≤ 0.001; **P ≤ 0.01; *P ≤ 0.05 and ns P >0.05. Three independent biological replicates were taken for each sample and lipids were extracted from 400 heads or 250 pairs of ovary or testis for each biological replicate.

**Fig.S2. CPES enzymatic activity is required for spermatogenesis and accumulation of ceramide is not responsible for cpes mutant phenotypes.** (A, C and E) Live testis squash preparation followed by phase contrast microscopy of round spermatids, (B, D and F) are immunofluorescence images of early elongating spermatids from samples corresponding to A, C and E respectively. Immunostaining was performed with beta tubulin primary antibody and Alexa Fluor 568 conjugated secondary antibody (Red). Nuclei are stained with DAPI (Blue) (A and B), the UAS-CPES active site mutant (GX_3_DX_3_D to GX_3_AX_3_A) was expressed using bam-Gal4 in cpes mutant background. (C and D), cpes; dcert^1^ double mutants. (E and F) The UAS CDase (neutral ceramidase) was expressed using bam-Gal4 driver in cpes mutant background.

**Fig.S3. Analysis of cell type specific transcriptomic signatures in cpes mutants:** (A) GSEA using top 50 cell type specific expressed gene sets and compared between cpes versus wild type (WT) and bam-Gal4>UAS CPES rescue (Rescue). Red bars indicate gene sets enriched in cpes mutants and blue bars indicate gene sets enriched in WT & Rescue. (B) enrichment plots for each of the cell type shows the distribution of individual genes in each set (vertical lines on red to blue parallel bar). Genes with positive enrichment score (ES) indicate that they are enriched in cpes mutants whereas negative enrichment score indicate that they are enriched in WT and Rescue. (C) Heatmap comparison of cell type specific enriched gene sets in w wild type (W1118), cpes and Rescue (bam-Gal4>UAS CPES rescue). Red indicates gene sets enriched in cpes mutants and blue indicate gene sets enriched in WT & Rescue.

**Fig.S4. Cell type specific expression pattern of various UAS and Gal4 constructs in Drosophila testis.** (A-) Whole mount Drosophila testis immunostained GFP with anti-GFP antibody (Green), DAPI stains DNA (Blue). (A) nanos-Gal4 (nos) drives the expression of pUASP-αTubulin-GFP, (B) bam-Gal4 drives the expression of pUASP-alphaTubulin-GFP, (C) chiffon-Gal4 drives the expression of alpha-Tubulin-GFP, (D) C587-Gal4 drives the expression of alpha-Tubulin-GFP, (E) nanos-Gal4 (nos) drives the expression of pUAST-EGFP, (F) bam-Gal4 drives the expression of pUAST-EGFP, (G) chiffon-Gal4 drives the expression of pUAST-EGFP, (H) C587-Gal4 drives the expression of alpha-Tubulin-GFP, (I) nanos-Gal4 (nos) drives the expression of pUAST-PLCδ-PH-EGFP, (J) bam-Gal4 drives the expression of pUAST-PLCδ-PH-EGFP, (K) chiffon-Gal4 drives the expression of pUAST-PLCδ-PH-EGFP, (L-N) Verification of various nanos-Gal4 drivers and their expression pattern. BDSC#4937 (used in this study) and BDSC#64227 specifically express transgene in GSC and early spermatogonia compared to BDSC#32563 which expresses more broadly.

**Fig.S5. Insect CPE synthases but not sphingomyelin synthases rescue spermatogenesis defects.** (A-C) Phase contrast microscopy of round spermatids where UAS hSMS1 (A), UAS Aedes CPES (B), and UAS Bombyx CPES (C) were ubiquitously expressed in cpes mutant background using tubulin Gal4. The scale bar is equivalent to 10 μm. (D) Measurement of cytokinetic defects in UAS-hSMS1, UAS-Aedes aegypti and UAS-Bombyx mori CPES rescue testes. 4 independent experiments for UAS-hSMS1 (round spermatid count, n=500), 2 independent experiments for UAS-Aedes aegypti (round spermatid count, n=1100) and 4 independent experiments for UAS-Bombyx CPES (round spermatid count, n=1900). (E-G) Confocal images of elongating spermatids immunostained with beta tubulin primary antibody and Alexa Fluor 568 secondary antibody (Red) and DAPI for DNA (Blue). The UAS hSMS1 (E), UAS Aedes CPES (F) and UAS Bombyx CPES (G). (H-K) Sphingolipid analysis of lipids extracted from testis of wild type, cpes mutant, and various rescues expressed in germ cells using vasa-Gal4. In contrast to Figs.1 and 2, here we used whole male organ (testis, accessory glands, collecting ducts and ejaculatory bulb) for lipid extraction. (H) The amounts of CPE and its subspecies in wild type, cpes mutant and various rescues are shown. (I) Ceramide and its subspecies are shown in picomoles per 100μg of carbon. (J) Hexosylceramide subspecies amounts are shown in bar diagram. (K) Amount of sphingomyelin (SM) and its subspecies are shown as picomoles per 100μg of carbon. Fluorescent blue green part of the bar represents SM species with net two double bonds in them. (L) Thin layer chromatography (TLC) of lipids extracted from 50 pairs of testes from each genotype. The position of SM on TLC is indicated by an arow. Dihydrosphingosine (DHSPH), Sphingosine (SPH), Sphingadiene (SPD) Saturated fatty acid (SFA), Monounsaturated fatty acid (MUFA) and polyunsaturated fatty acid (PUFA). Statistical significance was calculated using mean, standard deviation (SD) and N in Prism 8. The 2way ANOVA multiple comparison was used to calculate P values where ****P ≤ 0.0001;***P ≤ 0.001; **P ≤ 0.01; *P ≤ 0.05 and ns P >0.05. Three independent biological replicates were taken for each sample and lipids were extracted from 250 male organs for each biological replicate.

Fig.S6. GSEA pathway analysis indicates enrichment of spindle elongation and spindle elongation processes. (A) GSEA pathway analysis of RNA seq data from w^1118^, cpes mutant and bam-Gal4>UAS-CPES rescue testis. Three independent biological replicates were taken for each sample and RNA was extracted from 200 pairs of testes for each biological replicate. (B) Enrichment plot showing genes involved in spindle elongation are positively correlated in cpes mutants compared to w^1118^ and bam-Gal4>UAS-CPES rescue. (C-D) Heatmap shows spindle elongation genes are enriched in cpes mutants. (E) Enrichment plot showing genes involved in spindle organization are positively correlated in cpes mutants compared to w^1118^ and rescue. (F-G) heat maps show top 100 enriched genes in cpes mutants (H) Heatmap shows the top 50 genes enriched in wild type and rescue.

**Fig.S7. Cytological analysis of Rab11 and Rab7 mutants for meiotic cytokinesis defects.** (A) Measurement of cytokinetic defects in testes squash preparations of wild type samples, 7 independent experiments and total round spermatid count of about (n=6000). (B) cytokinesis defects in Rab11 dominant negative expressing gonads at 4^th^ day, 5 independent experiments and total round spermatid count of about (n=1500). Fraction of round spermatids showing nucleus to mitochondria ratio 2:1 (2nucleus), 3:1 (3nucleus), 4:1 (4 nucleus) and 4 irregularly sized nucleus with one mitochondrion are depicted. (c) Western blotting of protein extracts from control and Rab7 mutants using Rab7 and Tubulin (loading control) antibodies. Lanes 1-4 (control), lanes 5-8 (Rab7 mutants). (D) Fraction of cytokinetic defects in Rab7 null mutant pupal gonads (24-48 hours post pupation), 11 independent experiments and round spermatid count of about (n=1100). (E) Comparison of total cytokinetic defects between Rab11DN and Rab7 mutants (sum of all forms of cytokinetic defects including nuclear to mitochondria ratios 2:1, 3:1,4:1 and irregularly seized nucleus). Each dot in each graph represents an independent experiment (n) that consisted of 10 individual testis/gonads.

**Fig. S8 Sample processing for FIB-SEM**

A. Drosophila cysts were dissected onto a gridded cover slip, after fixation and embedment in 4% agarose. The cyst of interest is circled and is the same as the cyst shown in Fig. 7A. B. Higher magnification image of boxed area in A C. Light microscopy overview of full field before EM processing D. Same field after EM processing. The same pattern of dissected cysts is visible, suggesting spatial retention of sample. Trimmed area approximated with a dashed rectangle. E. Field of view (orthogonal to Fig. 7A) with doubly labeled cleavage furrow of interest circled. F. Sectioning at the microtome proceeded until furrow of interest was almost exposed (red circle). FIB-SEM imaging was executed in the volume indicated by rectangle, in direction of arrow.

## Notes

### Competing Interest Statement

The authors have declared no competing interest.

