## Supplementary figures and images for "Endosomes deliver ceramide phosphoethanolamine with unique acyl chain anchors to the cleavage furrow during male meiotic cytokinesis"

### Supplemental Figure 1

**Figure S1**
**Kunduri et al., 2022**

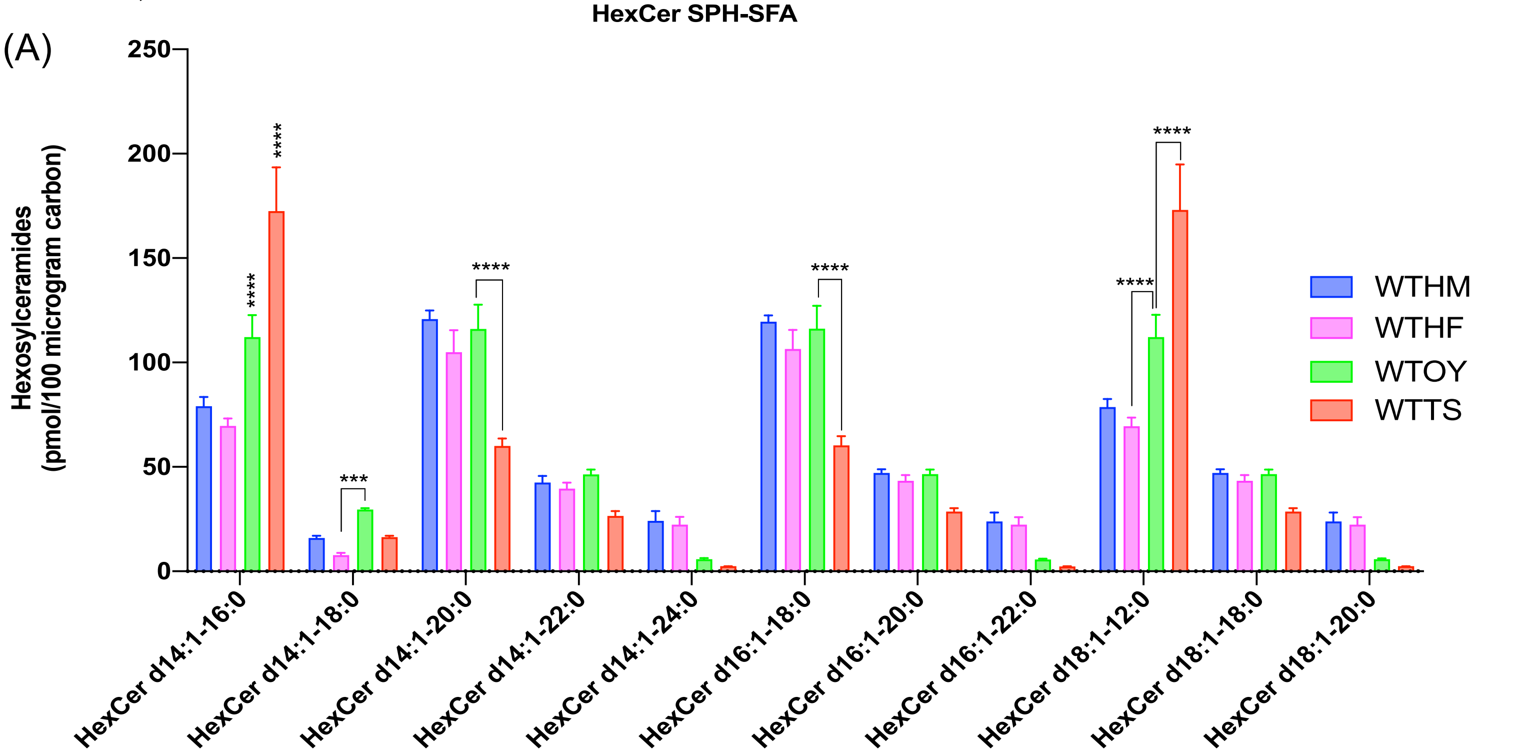


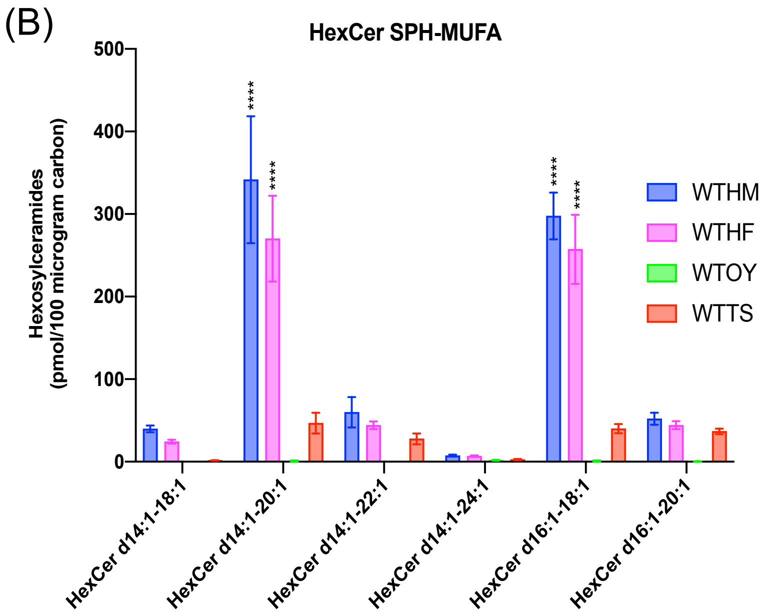


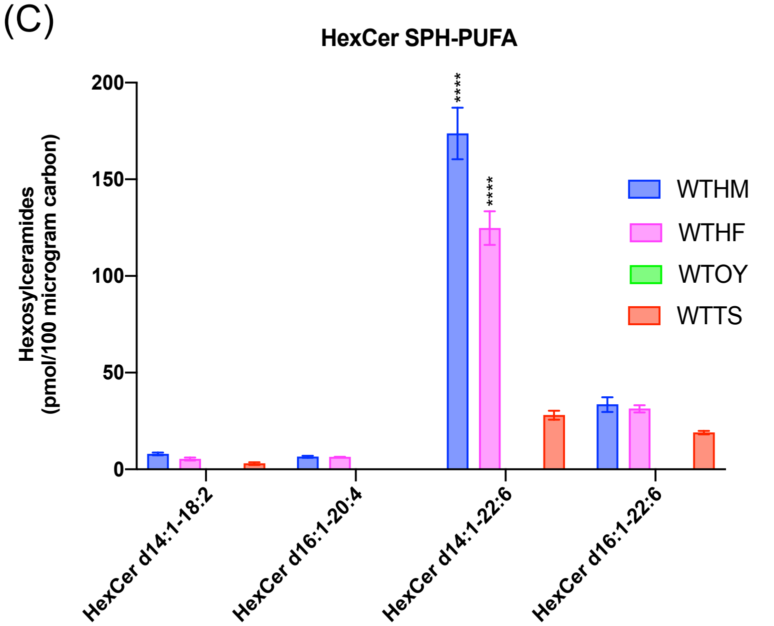


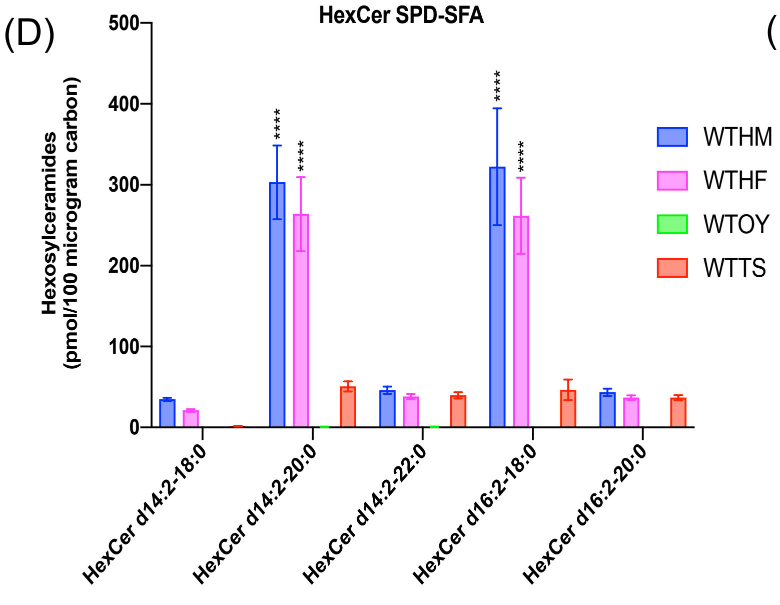


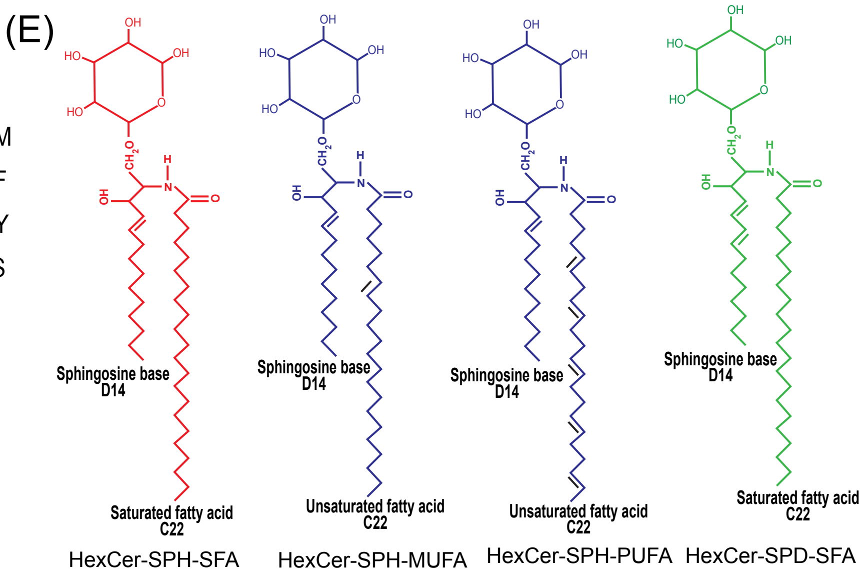

### Supplemental Figure 2

**Figure S2**
**Kunduri et al., 2022**

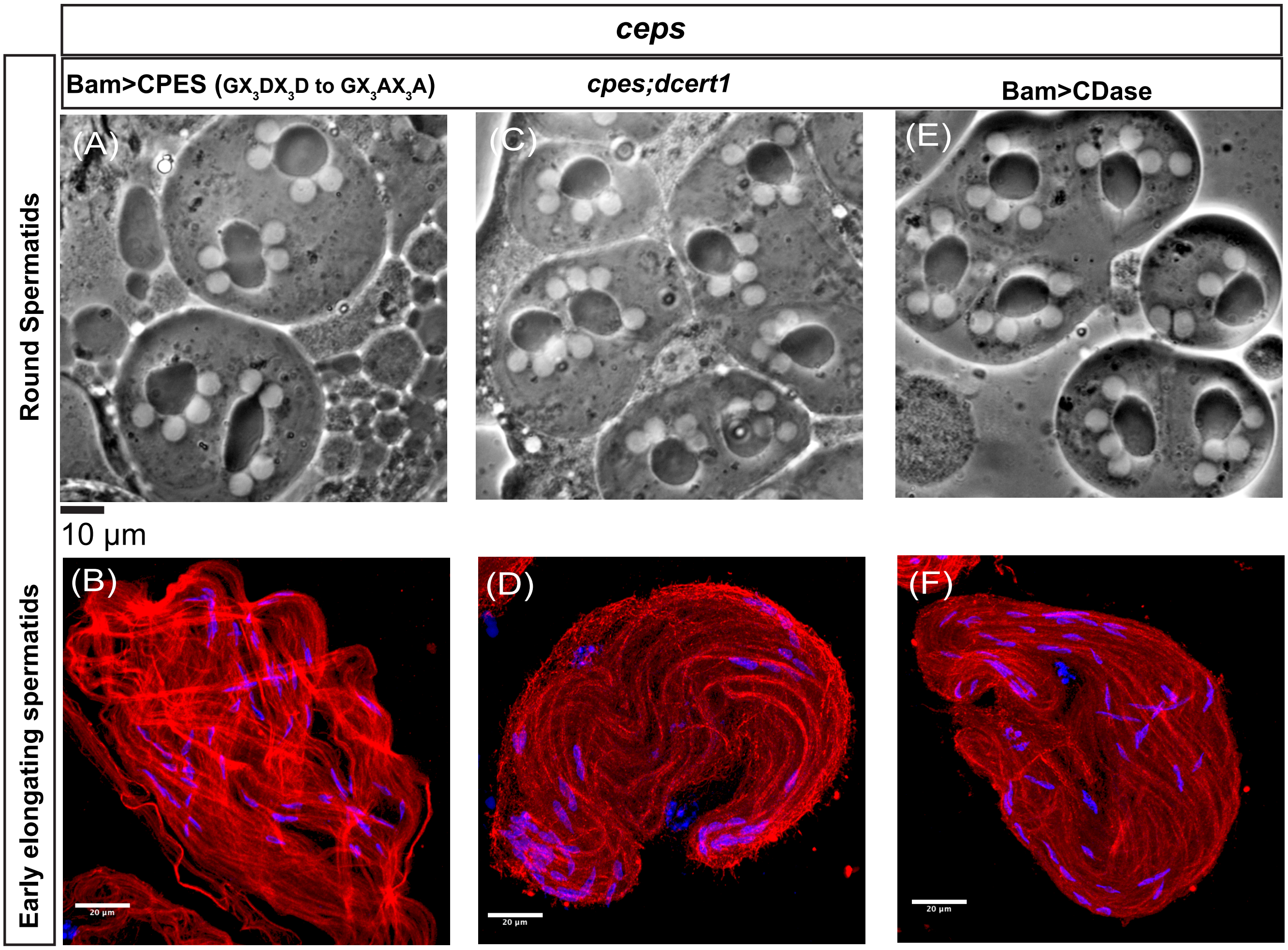

### Supplemental Figure 3

**Figure S3**
**Kunduri et al., 2022**

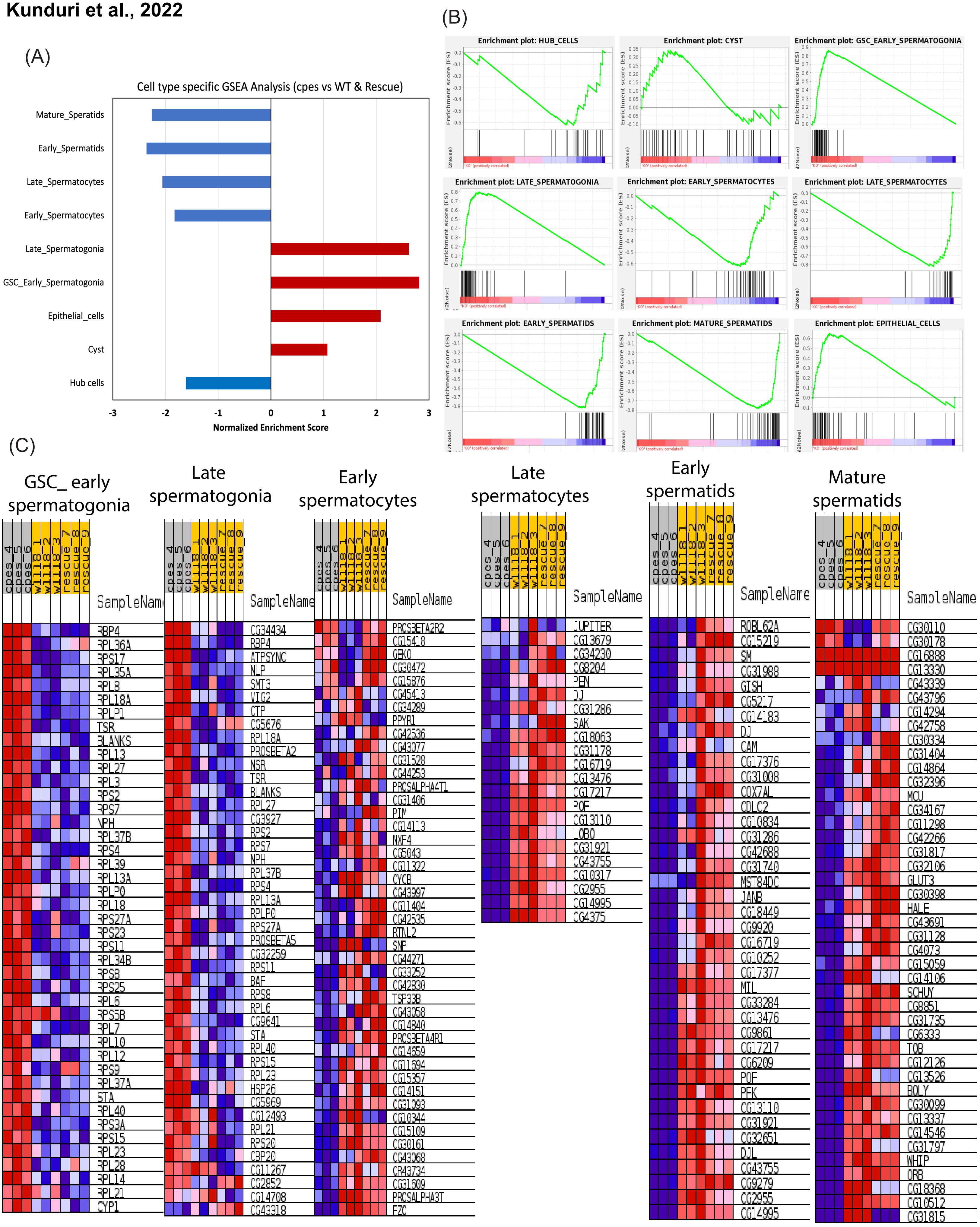

### Supplemental Figure 4

Figure S4
Kunduri et. al., 2022

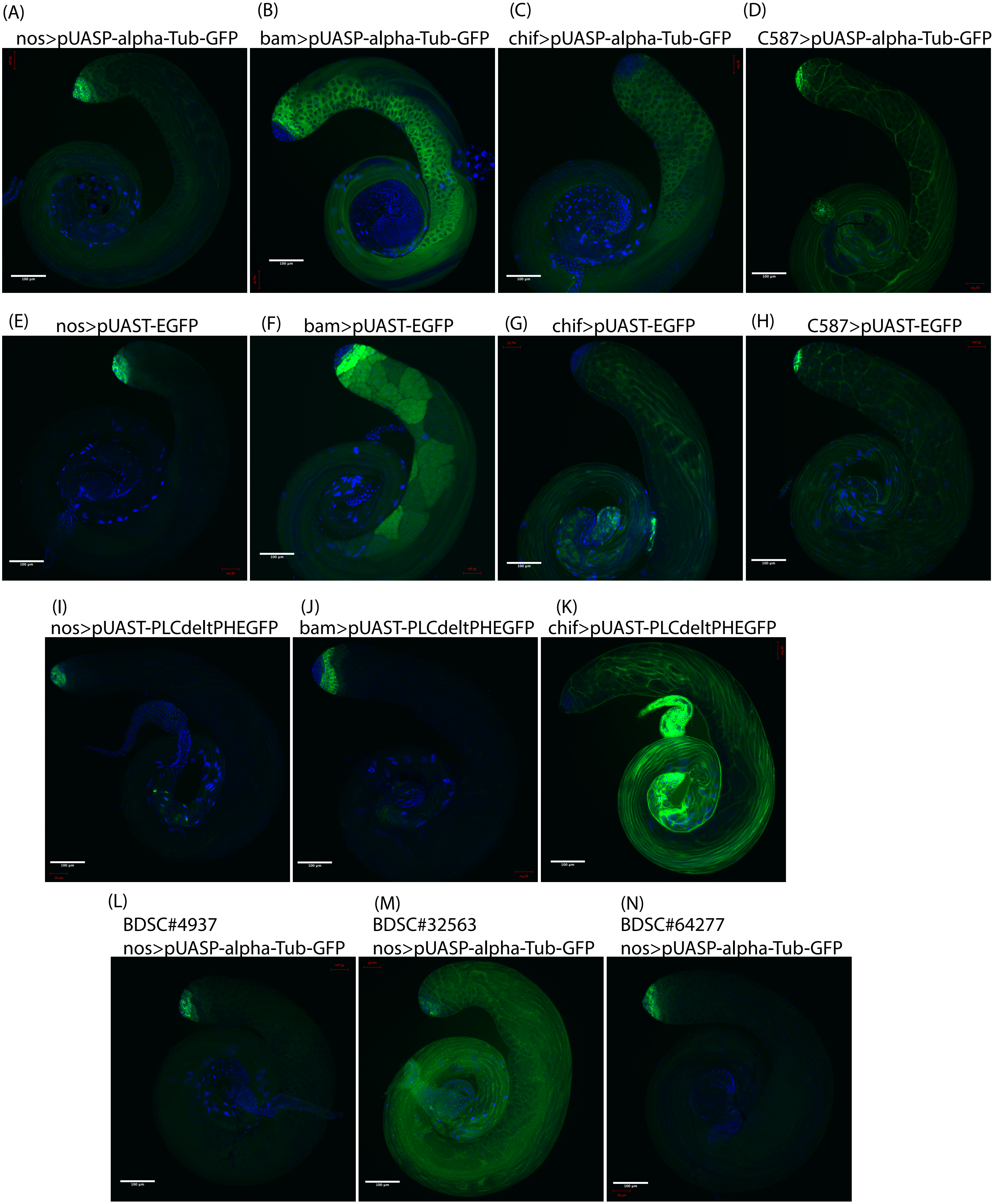

### Supplemental Figure 5

**Figure S5**
**Kunduri et al., 2022**

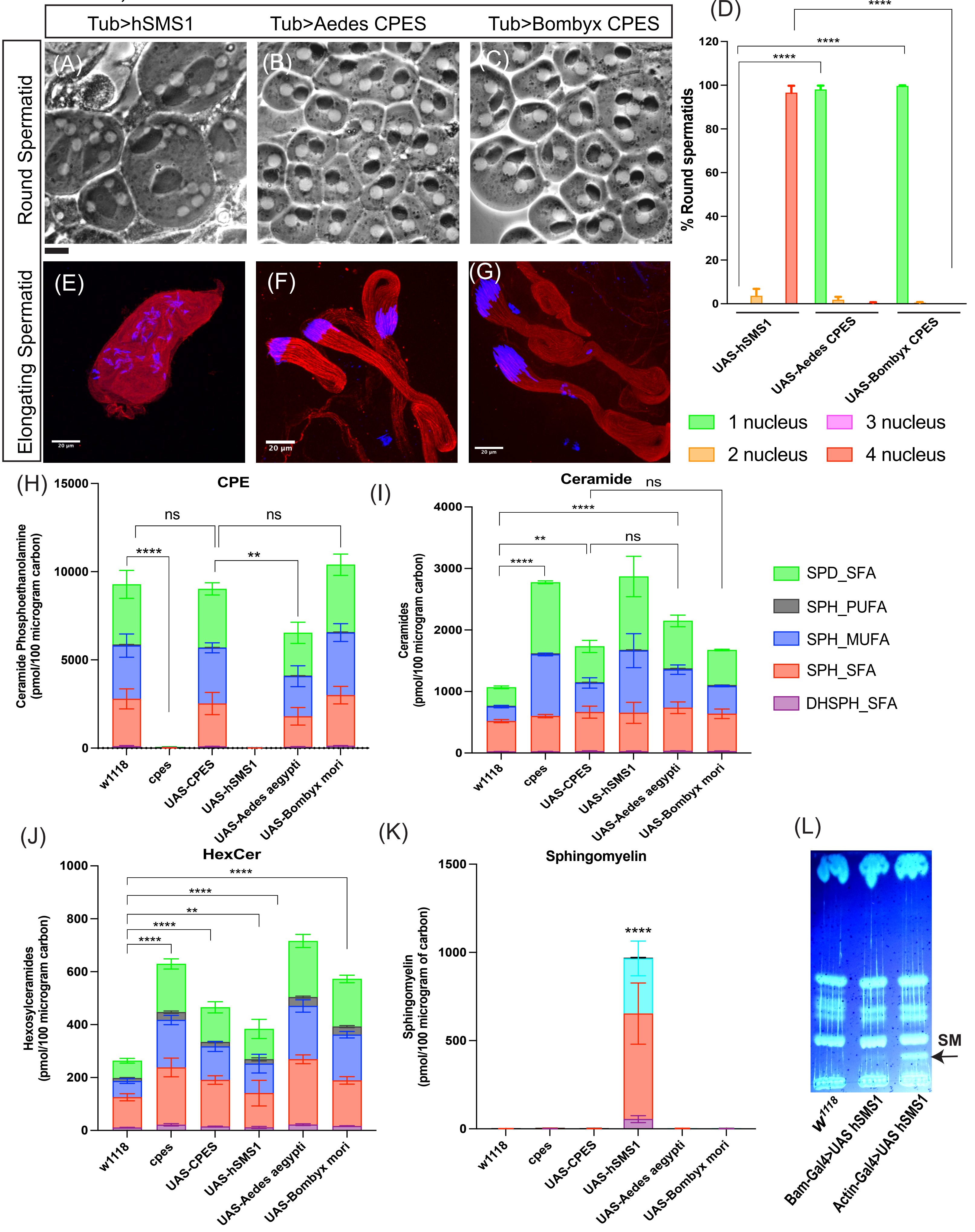

### Supplemental Figure 6

**Figure S6**
**Kunduri et al., 2022**

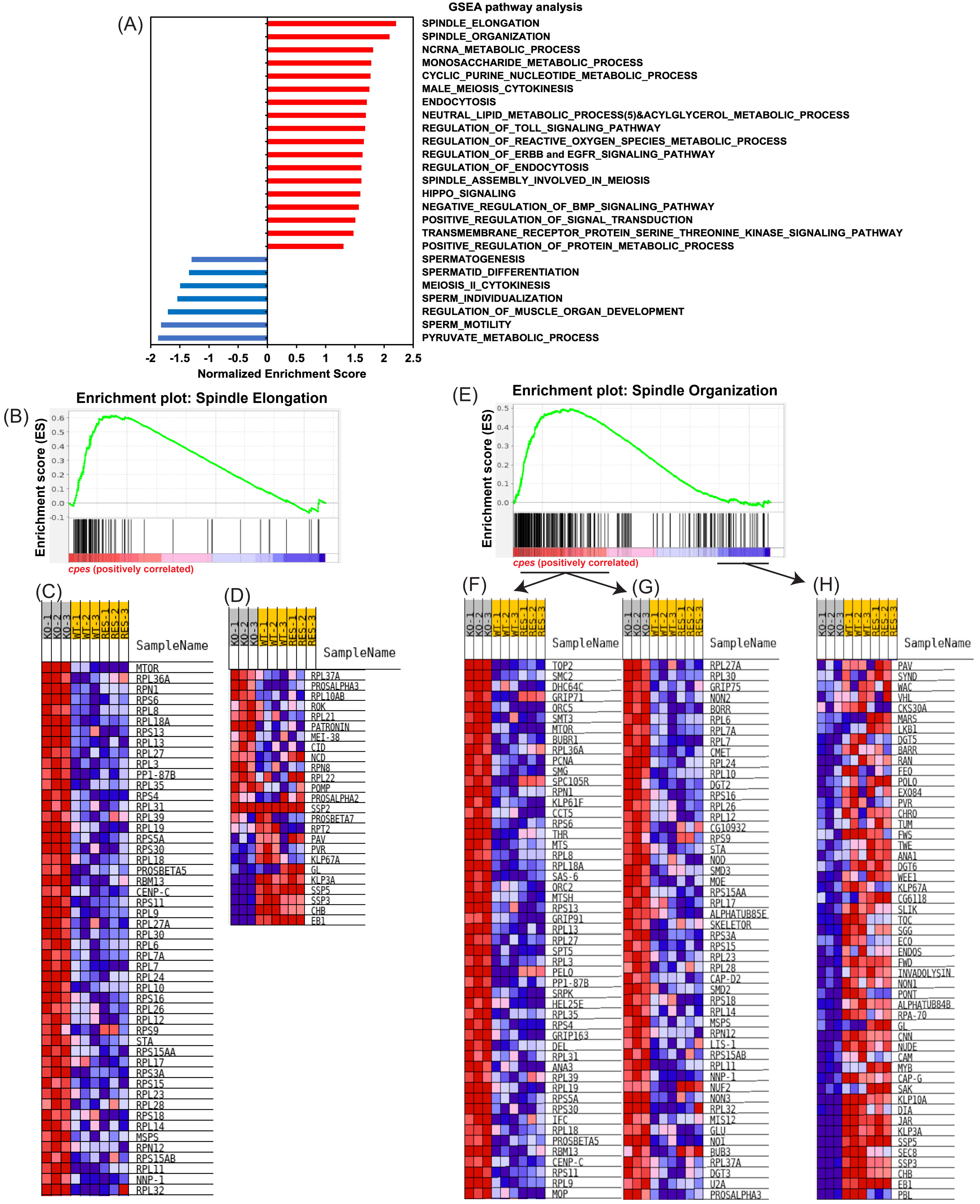

### Supplemental Figure 7

**Figure S7**
**Kunduri et al., 2022**

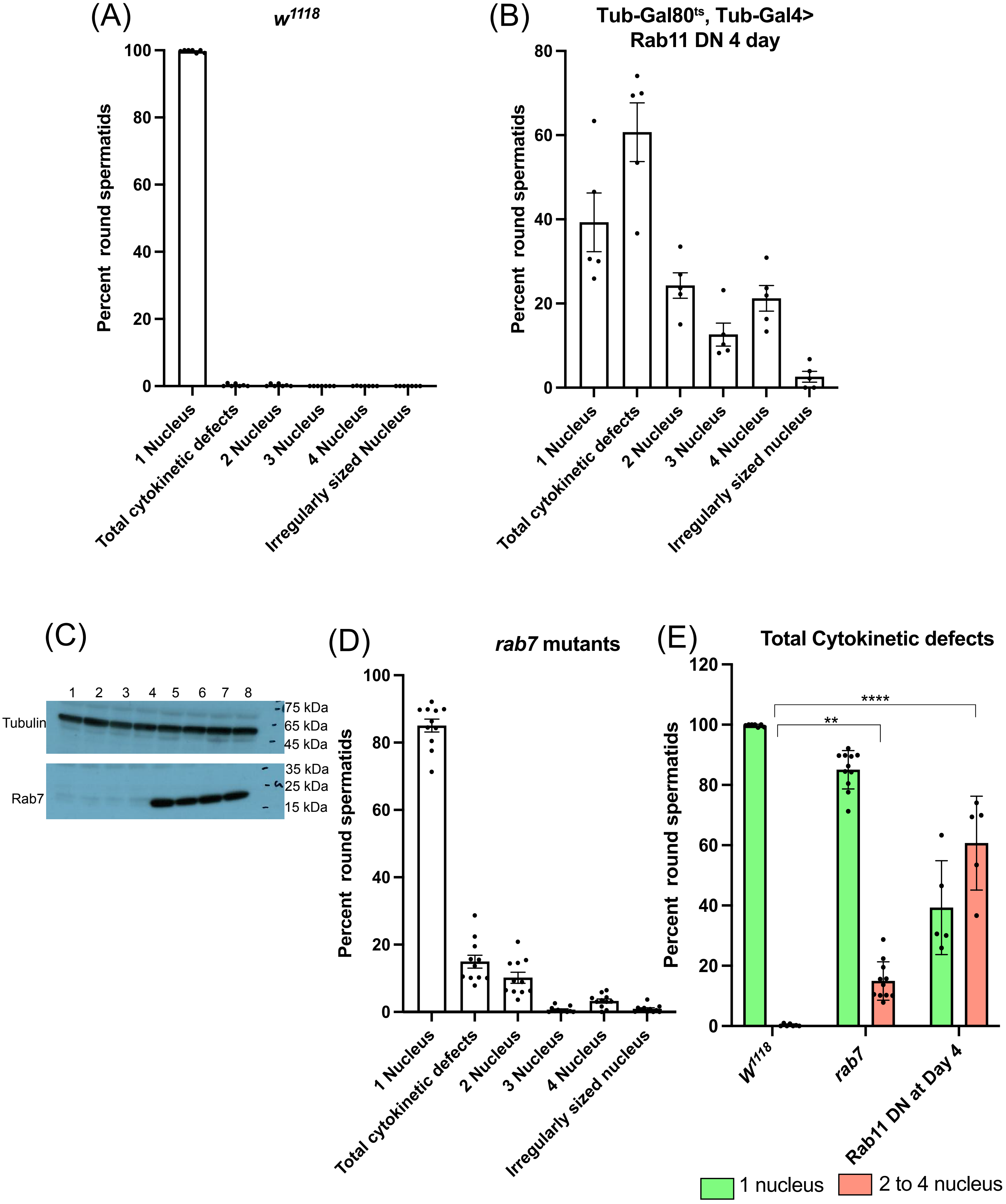

### Supplemental Figure 8

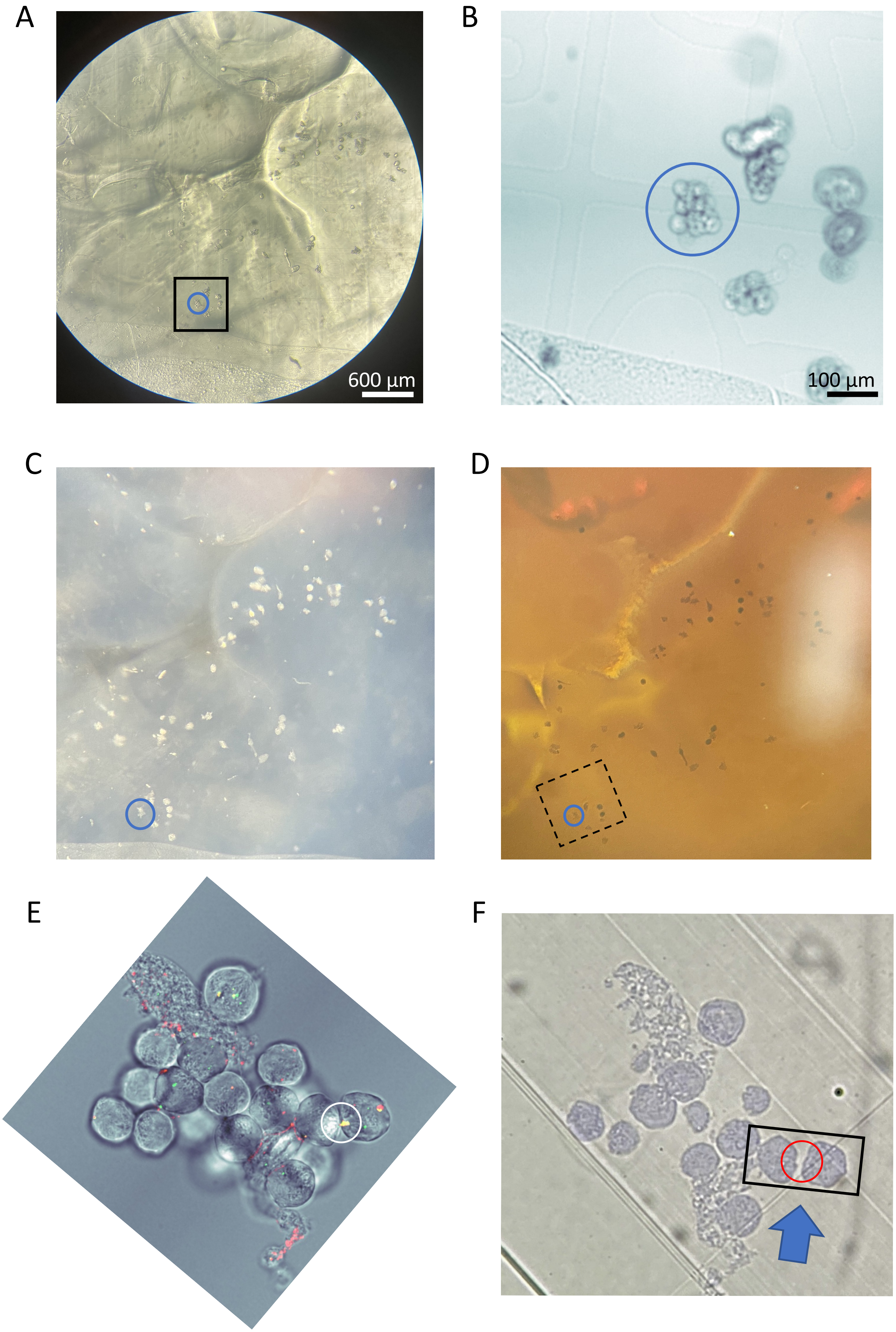
