## Supplemental Table S1 for "Endosomes deliver ceramide phosphoethanolamine with unique acyl chain anchors to the cleavage furrow during male meiotic cytokinesis"

| Name | Sequence (5’-3’) | Description |
| --- | --- | --- |
| Aedes FW | ATAgAATTCAAAATggTggggCCCAgTTCgCAAATC | To PCR amplify Aedes aegypti CPES ORF and subclone into PUAST vector; EcoRI site is underlined. |
| Aedes Rev | CCCTCTAgATTACgTAgAATCgAgACCgAggAgAgggTTAgggATAggCTTACCACCCAACgATgTATTgTTACCAgT | To PCR amplify Aedes aegypti CPES ORF with V5 tag at C-terminus and subclone into PUAST vector; XbaI site is underlined. |
| B mori FW | ATAgAATTCAAAATgTggCCACCATCgCAAgCCAgC | To PCR amplify Bombyx mori CPES ORF and subclone into PUAST vector; EcoRI site is underlined. |
| B mori Rev | CCCTCTAgATTACgTAgAATCgAgACCgAggAgAgggTTAgggATAggCTTACCACCTCTACCATCAACggAggTTTTAAC | To PCR amplify Bombyx mori CPES with V5 tag at C-terminus and subclone into PUAST vector; XbaI site is underlined. |
| T urticae FW | ATAgAATTCAAAATggCCACTTTTCTggCTgAAC | To PCR amplify Bombyx mori CPES ORF and subclone into PUAST vector; EcoRI site is underlined. |
| T urticae Rev | CCCTCTAgATTACgTAgAATCgAgACCgAggAgAgggTTAgggATAggCTTACCACCgTAATAAgAgAgTACAgAAgAAAC | To PCR amplify Bombyx mori CPES ORF with V5 tag at C-terminus and subclone into PUAST vector; XbaI site is underlined. |
| CPES active site FW | CCTCCggCTACTACgTAgCCggCCTgTgCgCCggACTCggCTgC | To replace active site Aspertates (D214A and D218A) with Alanine using site directed mutagenesis method. |
| CPES active site Rev | gCAgCCgAgTCCggCgCACAggCCggCTACgTAgTAgCCggAgg | To replace active site Aspertates with Alanine (D214A and D218A) using site directed mutagenesis method. |
| CPES FW | ACAAgATCTATgATCggACCCAgTTCgCAg | To PCR amplify CPES ORF with active site mutation and V5 tag at the C-terminus and subclone into pUAST vector. BglII site is underlined. |
| CPES V5 Rev | ACACTCgAgTCACgTAgAATCgAgACCgAg | To PCR amplify CPES ORF with active site mutation and V5 tag at the C-terminus and subclone into pUAST vector. XhoI site is underlined. |
| PlyA2Fw | CCCCATATggCCTACgCCCAgTgggTC | To PCR amplify PlyA2 ORF and subclone into pET24a vector for protein expression and purification. NdeI site is underlined. |
| mCherry Rev | CCCCTCgAgCTTgTACAgCTCgTCCATg | To PCR amplify PlyA2-mCherry and subclone into pET24a vector for protein expression and purification. XhoI site is underlined. |

Table S1
